# Habitat fragmentation controls bacterial community composition outcomes

**DOI:** 10.64898/2026.02.04.703711

**Authors:** Maxime Batsch, Anna Puzyrko, Isaline Guex, Jan Roelof van der Meer

## Abstract

Natural habitats of bacteria are often highly structured in space, but how this affects their interactions and community composition outcomes is poorly understood. To address this gap, we studied the development of a synthetic soil community of 21 bacterial strains under a gradient of habitat fragmentation on single or mixed-substrate conditions. Fragmentation was produced by random microfluidic seeding of the mixed inoculum into culture medium droplets of variable volume (30-150 pL), targeting mean starting densities between 1 and 27 founder cells comprising 1-2, 7 or 14 taxa per droplet. Across all droplets for each of the fragmented states, globally different community compositions arose, suggesting variable individual taxa strategies and adaptation to differences in habitat fragmentation. In comparison to the same mixed inoculum growing under the same nutrient conditions in bulk culture, the more fragmented conditions allowed higher cell density and diversity, with less overgrowth by a few strains. Community evenness was highest when inocula were dispersed into individual taxon (1-2 taxa) or low-order combinations (7 taxa per droplet), but individual dispersion reduced the productivity of the meta-community across droplets. Monod-type simulations suggested that most community composition could be explained by the inherent individual kinetic properties and dispersal constraint effects of the fragmented habitat, with smaller effects from emerging interspecific interactions. Overall, our results show the strong effect of habitat structure on community outcomes, with higher fragmentation states favoring more even species coexistence.

**TEASER:** Habitat fragmentation results in the formation of locally isolated, smaller communities with varying richness and interactions, which altogether contribute positively to the maintenance of prokaryotic species diversity.

## Introduction

Understanding how microbial communities assemble and what allows so many different taxa to co-exist in nature is one of the key questions in microbial ecology. A growing body of literature emphasises the central role in community assembly of nutritional niche preference differences among taxa^1–3^, as well as biomolecular interactions (*e.g.*, appendices, bioweaponry)^4–9^. Taxa compete depending on their nutritional niche preferences and on the (primary) resource availability, resulting in dynamic patterns of metabolites being produced, leaked, and reused, all of which control individual population growth and emergent community behaviour.

Most experimental studies acknowledge that resource availability and interspecific interactions are tightly coupled to emergent community structure, but few have specifically addressed the role of spatial niches on community development^10^. As a matter of fact, most natural microbial habitats are fragmented or disconnected at a size relevant for microbial cells or colonies (*i.e.*, 0.2–100 µm), because of the existence of microstructures, cavities^11–13^, water availability, particles or other^9,14^. Typical examples include the water-filled cavities and micropores of the soil matrix, or microdroplets formed on leaf surfaces^15^. Floating particles in aqueous systems can also be considered disconnected habitats. Pores, particles, or droplets contain, adsorb, or confine low numbers of cells, segregating potentially highly diverse communities into collections of smaller sub-communities with fewer taxa because of spatial constraints and imposing local compositional heterogeneities^16,17^. One would expect that such spatial discontinuities have a direct effect in limiting the number of cells and taxa being present in the same site, or even isolating certain taxa from others. These heterogeneities will then at least temporarily restrict the possible interactions between all taxa in the community to only those cells of the taxa present in close vicinity. This will then define the types of metabolic exchanges and biological interactions, such as the effects of secreted secondary metabolites^8,18^ or contact-mediated killing processes^19^. For example, spatial micro-discontinuities were shown to stabilize the coexistence of bacterial populations in a predator-prey relationship^20,21^. Furthermore, micro-fragmented habitats were shown to favor higher than expected proliferation of poorly competitive strains because of phenotypic variation among founder cells^22^. Recent work has also suggested that the degree of habitat fragmentation influences microbial diversity depending on the prevailing sign (positive or negative) of interspecific interactions within a community of *E. coli* strains^23^.

Microbial communities in fragmented habitats may thus be considered as a sum of parallel micro-scale communities, whose assembly processes involve locally different and dynamic combinations of subsets from the overall available taxa composition. Experimental studies on community assembly processes so far have frequently been carried out in homogenised (all taxa present) and completely mixed liquid culture systems, leading to a loss of spatial micro-fragmentation^2,4,5,24^. Homogenisation removes the variation in emerging interspecific interactions under fragmentation, which might both amplify or reduce effects on individual taxa growth in community assembly. We hypothesize that fragmentation plays a dominant role because it controls local species assemblages by seeding bottleneck effects and restructures the types and magnitudes of interactions that can emerge. This, we expect, will affect community compositional and diversity outcomes.

To address our hypothesis, we took advantage of a system of culturing in picoliter water-in-oil droplet emulsions, which results in numerous spatially separated habitats for microbial growth with volumes relevant to natural scale fragmentation (*e.g.*, 40-150 pL)^24,25^. Droplets are produced by microfluidics, and bacterial cells are Poisson-randomly encapsulated within the droplets from a starting suspension, which can be precisely designed and mixed from individual pure cultures to the intended strain relative abundances^25,26^. Our target community here consisted of 21 cultured soil bacterial isolates (a soil SynCom defined in Ref^27^), grown individually and then mixed at the start to the same relative abundances. To test the effect of habitat fragmentation and the interaction order (*i.e.*, the number of locally available species), we diluted the starting SynCom to different cell densities under increasing droplet volumes (Fig. 1a). This resulted in SynCom subset combinations of 1–2 cells and species per droplet (43– µm volume, called here *isolation* or I), 7 cells and species (43–µm volume droplets, *low-order* or *L*, range 1–15 cells and 1–11 species), 27 cells (range 15–40) with 14 species (range 8–19) per droplet (67–µm droplets, *high-order* or *H*). In comparison, we also incubated the SynComs without fragmentation in bulk culture (200 µl volume with 21 species, called *F* for *full*, Fig. 1a). In addition, two medium conditions were imposed: one with a single carbon and energy substrate (succinate), for which we expected substrate competition to become dominating, and the second with a substrate mixture (an aqueous extract of soil organic matter, or *SE*), which we expected to present different nutritional niches for the SynCom strains, facilitating their coexistence. Droplet emulsions and bulk liquid cultures were incubated under aerobic conditions for 7 days, and total community growth was followed by flow cytometry at different incubation time points. Growth in individual droplets was further assessed from changes in bright-field contrast scatter in microscopy images. Relative strain abundances were determined from their unique signatures present in amplified and sequenced 16S rRNA gene V3-V4 regions on sample-purified total DNA. Finally, to corroborate community development under fragmented and bulk conditions, and estimate the contribution of both inherent individual strain growth kinetics as well as potential interspecific interactions, we adapted a Monod growth kinetic model^28^ to droplet emulsions and Poisson-randomised starting species compositions. Our results demonstrate the strong importance of habitat fragmentation on emerging community composition outcomes, regardless of the imposed substrate conditions.

**Figure 1:**
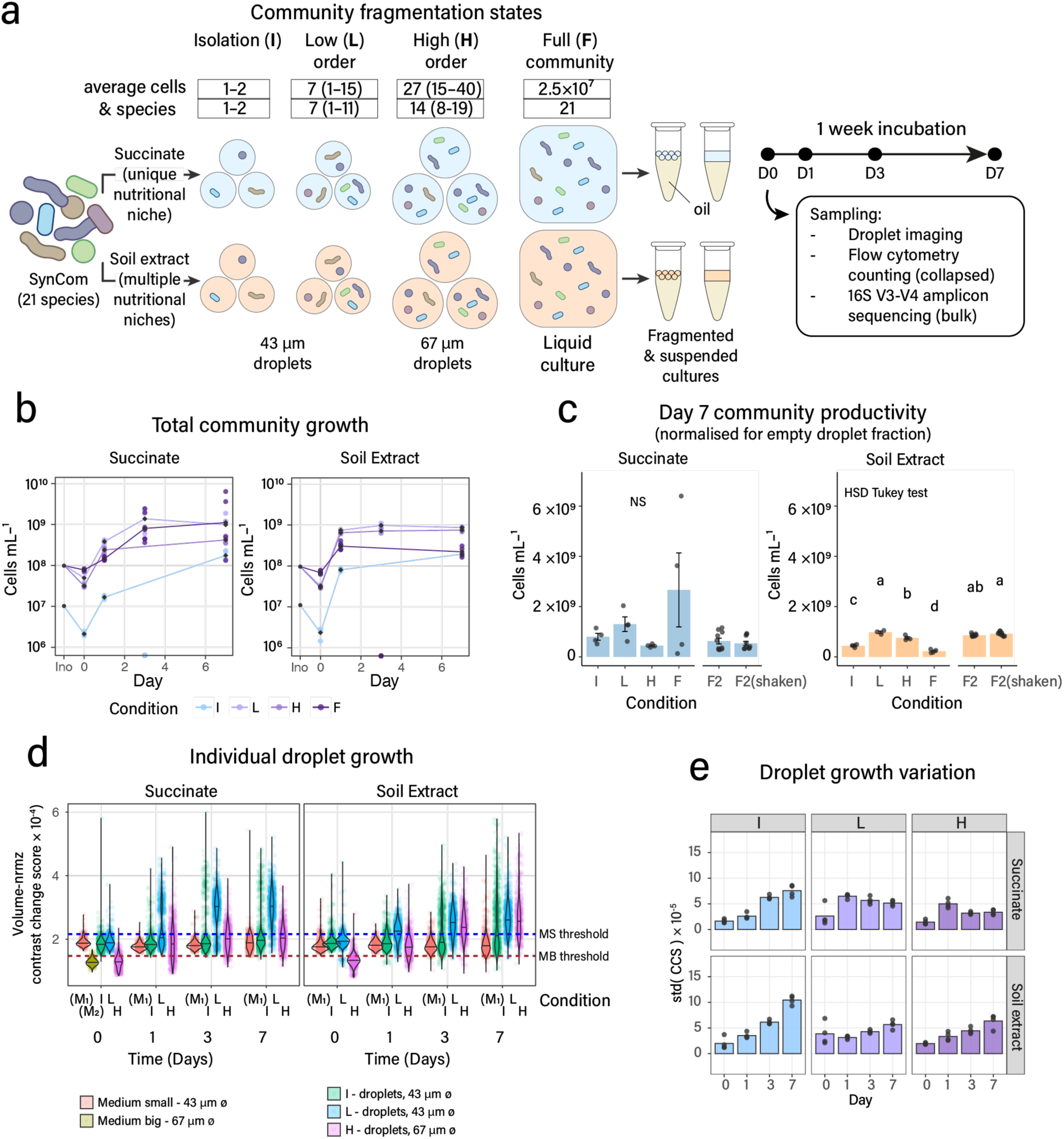
Total and per droplet SynCom growth varies with habitat fragmentation and resource context. **a** Experimental design to vary habitat fragmentation states and local species order (I, isolated taxa; L, low order; H, high order, and F, full community bulk suspension), with two media conditions. Boxes indicate the starting number (and range) of cells and taxa per droplet or bulk condition. D0, D1, D3, D7; sampling days and associated measurements. **b** Total community growth under the various conditions measured by flow cytometry. Lines connect the means per condition (*n* = 4 replicates) and time point. Circles show individual replicate values. Note that the D3 samples for some of the conditions were lost. Ino, community size measured of the inoculum before droplet generation. **c** Comparison of total community sizes after 7 days of growth. F2 denotes an independent repetition of the bulk condition to test the effect of sample resuspension. Bars show mean community sizes with circles indicating individual replicates and the error bars depicting standard deviations. Letters indicate statistically significant differences between conditions at a threshold p < 0.05 from an HSD Tukey test. ‘NS’, not significant. **d** Individual droplet growth as contrast change score (CCS) from bright-field microscopy. Violin plots show circles depicting individual droplet measurements, colored by condition as per the legend below. I, L, and H refer to the fragmentation condition as in (a). M1, M2, medium background in empty droplets of the same volume. Dotted lines indicate the 95^th^ percentile of the medium droplet background CCS-value, above which droplets were considered to contain cells. **e** Variation of individual droplet growth displayed as the standard deviation of droplet-CCS (including empty droplets) per condition and time point. Bars show the mean of four replicate values, with circles displaying individual replicate values.

## Materials and methods

### SynCom assembly and culturing

Individual soil SynCom strains were streaked on R2A agar plates from glycerol stocks stored at -70°C. After up to two weeks (depending on the strain) of incubation at room temperature, surface-grown strain biomass was resuspended in 500 µl of Soil Buffer (per L: 0.6 g of MgSO_4_·7H_2_O, 0.1 g of CaCl_2_ and 1.8 mL of 5 x M9 minimal salts solution [BD Biosciences]), which was transferred into 2-mL Eppendorf tubes. The tubes were then loaded on a multi-tube holder (SI-V525 Vertical, Scientific Industries), which was placed on a Vortex-genie 2 (Scientific Industries) for at least 20 min at speed 8 to disperse the cell biomass. The turbidity (OD_600_) of the resulting suspension was measured and adjusted with soil buffer to OD_600_=1. Aliquots of 400 µL of the diluted suspension of each of the strains were then mixed in a 50-mL Falcon tube, resulting in a total volume of 8.4 mL SynCom cell suspension.

Separate triplicate mono-species suspensions with a turbidity of ca. OD_600_ = 1 were fixed (sodium azide, final concentration 4 g/L) and stained with SYBR Green I (Thermofisher, following supplier’s instructions), and counted by flow cytometry to derive the ratio between OD and cell numbers (Supplementary Fig. 1). This was used to calculate the relative strain abundances in the starting mixed inoculum.

The final 21-species mixture was divided over two 15 mL tubes (each with 4.2 mL), which were centrifuged for 8 minutes at 4000 × *g* to collect the cells, after which the supernatant was discarded, and the cell pellet was carefully resuspended in 4 mL of soil buffer. After another round of centrifugation, the cell pellet was then either resuspended in 4 mL of 21C minimal medium + 10 mM succinate^22^, in 21C medium without carbon substrate (to count the initial cell per droplet distribution, see below), or in 4 mL of sterile aqueous soil extract (SE, preparation as described by Causevic et al. 2022^27^). In preliminary experiments, we also included 40 µM Na_2_-resazurin in the growth media to follow per-droplet cell respiration from resazurin reduction into the red fluorescent Resorufin^29^ and correlate this to cell growth measurements from bright-field contrast scatter (see below). Both SynCom mixtures were then diluted further in their respective media to a final OD_600_= 0.035 (for the L, H and F-conditions), or 0.0035 (for the I-condition). OD_600_= 0.035 corresponded to a starting inoculum of approximately 10^8^ cells/mL and allowed to produce on average 7 and 14 starting taxa per droplets in L- and H-conditions, respectively. OD_600_= 0.0035 corresponded to a starting cell density of approximately 10^7^ cells/mL and was used to produce droplets with 1-2 taxa in the I-condition. 300 µl of either of the mixed suspensions was used to generate droplet emulsions cultures using microfluidics (I, L, and H-conditions; see below) or added to Eppendorf tubes directly (F-condition, 300 µl per tube). Four replicates were produced for each condition. Droplet emulsions and bulk liquid cultures were similarly incubated without shaking at room temperature (21±2 °C) for a week in the dark with intermittent sampling (day 0 post-encapsulation, day 1, day 3 and day 7).

To extract individual growth kinetic parameters, monoculture strain suspensions at OD=1 were diluted 1:30 (5 µl in 145 µl) in triplicates into either 21C minimal medium with 10 mM succinate or SE, and dispatched in wells of a 96-well plate. The plate was then incubated for 7 days under continuous shaking (double orbital, 282 cpm, slow orbital speed) at 21 °C in a plate reader (BioTek, Synergy H1) during which the culture turbidities (OD_600_) were measured every 30 minutes. To limit evaporation, a line of H_2_O-filled wells (140 µl) surrounded the cultivation wells. Culture turbidity reads (OD_600_) were ln-transformed, and the manual beginning and end of the exponential phase for each strain were defined on an interactive plot (MATLAB R2021b), from which the ln-linear slope was calculated. The mean from the triplicate values was then taken as the µ_max_ of that strain on the substrate.

### Cell droplet encapsulation

SynCom starting suspensions of either OD=0.035 or OD=0.0035 were loaded into four separate 1 mL syringes (Omnifix 1 mL, U-100 Insulin), which were connected in parallel to a single chip carrying four droplet generator devices (40 µm × 40 µm × 40 µm cross junction, design from Duarte et al.^30^, custom-produced by Wunderlichip GmbH, Switzerland). Dispensing tips (20 ga, Luer), PTFE-tubing (OD 1.6 mm (1/16’), ID 0.8 mm (1/32’), Adtech Polymer Engineering™ BIOBLOCK/14), and stainless steel couplers (20 ga x 15 mm Instech Laboratories stainless steel catheter coupler SC20/15) were used for the junctions between syringes, PTFE-tubings and chip inlets. HFE 7500 fluorinated oil + 2% surfactant (008-fluorosurfactant, Ran Biotechnologies) was loaded in a separate 1 mL syringe and connected to the oil inlets on the chip. Syringes were mounted on multisyringe pumps (44/22 Mec, Harvard Apparatus) to inject the SynCom suspension at flow rates of 8 (for the 43 µm ø droplets) or 20 µL/min (for the 67 µm ø droplets), and the oil-surfactant mixture at 20 or 10 µl/min, respectively. The OD 0.0035 mixture produced droplets of 43-µm ø; whereas OD 0.035 was used to produce both 43- and 67-µm ø droplets (Fig. 1a; ranges see main text). The chip outflow was connected to 1.5 mL Eppendorf tubes to collect the generated droplets during 25 min (low flow rate) or 13 min (high flow rates) to obtain equivalent droplet emulsion volumes in the tubes. Medium-only emulsions of either medium (21C with succinate or SE) were prepared for both 43- or 67–µm droplet diameters to threshold the contrast change score-calculation for detecting cells (see below).

### Quantification of the starting SynCom cell distribution in droplets

To quantify the starting cell number distribution in culture droplets, one ml of the SynCom mixture at OD_600_ = 0.035 in 21C medium without carbon substrate was stained with 20 µl of SYBR Green I (100-fold diluted in DMSO from the purchased stock solution, Thermofisher) for 15 min, protected from light, before encapsulation in 43-µm ø droplets (Supplementary Fig. 2). Droplets with stained cells were transferred into a droplet chamber chip with a low 10-µm ceiling (design from Taylor et al. 2022^31^, chip preparation as previously described^22^) for imaging of individual cells by epifluorescence microscopy.

### Droplet imaging

To image growth in individual droplets, a 1 µL-droplet emulsion was subsampled at the defined incubation timepoints and transferred into a Countess chamber slide (Invitrogen C10228) containing 5 µL of HFE 7500 fluorinated oil. After pipetting the emulsion, another 5 µL of oil was added, and droplets were allowed to disperse for 1-2 min into a monolayer in the slide. The slide was then imaged on a Nikon Ti-2000 inverted epifluorescence microscope equipped with a Flash4 Hamamatsu camera and a 20x objective (Nikon CFI S Plan Fluor ELWD 20XC MRH08230). Images were taken in bright field (BF, exposure time = 25 ms), and exported as 16-bit TIF files. In case of quantifying Resorufin fluorescence, we additionally imaged droplets in the RFP-channel (red fluorescence, LED intensity = 100%, exposure time = 250 ms), whereas for quantifying SYBR green I-stained cells we imaged droplets under green fluorescence (GFP-channel, LED intensity = 100%, exposure time = 250 ms).

### Droplet and cell segmentation

Droplet images were processed and analysed with a MATLAB (R2021b, MathWorks Inc.) custom script. Individual droplet boundaries were identified using the *imfindcircles* function, and cell biomass inside droplet areas was estimated from the contrast change score (CCS), following the approach described by Kehe et al. 2019^29^. In short, the CCS is the sum of the standard deviations of each pixel grayscale value calculated within a surrounding kernel of 7×7 pixels within the droplet boundary. We used the 95^th^ percentile of CCS of empty droplets (either 43- or 67-µm ø of either medium) as the threshold above which we assume cell biomass can be detected (Supplementary Fig. 3, Fig. 1d). In a separate experiment we determined the correlation between CCS and resorufin fluorescence intensity (from resazurine respiration; as the sum of all pixel intensities in the RFP channel within the droplet boundary; Supplementary Fig. 4). SYBR Green I-stained cells were segmented within droplet boundaries using the segmentation process as described previously^22^.

### Flow cytometry counting of community sizes

Fixed and stained cells were counted by flow cytometry to quantify community growth in emulsions and mixed liquid cultures over time. In the case of droplet emulsions, we sampled and transferred a 40 µl emulsion aliquot to a 200-µl PCR tube, which was mixed with 40 µl of 1H,1H,2H,2H-perfluoro-1-octanol (PFO, diluted 4 times in HFE 7500 fluorinated oil). PFO-droplet emulsions were briefly (5”) vortexed and centrifuged on a mini-table centrifuge (C1301B-230V, Labnet International) for 10“ to separate the aqueous (bottom) and the PFO-oil phases (top). The PFO-oil phase was carefully removed, and the aqueous phase was transferred to a 2-by-2 cm parafilm square to remove residual oil (which sticks to the parafilm). The aqueous phase was pipetted from the parafilm (recovering about 20-25 µl from 40 µl of emulsion) into clean PCR tubes. 10 µl of this aqueous suspension was then counted by flow cytometry, and the remaining suspension was frozen at –20°C until DNA extraction. For the full mixed liquid culture condition, 20 µL was sampled, of which 10 µl was used for flow cytometry, and 10 µl was again frozen for DNA extraction.

Samples for staining were further diluted 1:100 in sterile PBS and mixed (50 µl) with 50 µl of 8 g/L sodium azide (NaN_3_) to fix the cells for 15 minutes at room temperature. Cells were then stained by the addition of 1 µl of SYBR green I (10,000 times pre-diluted in DMSO from the commercial stock) and incubated for 15 min in the dark, after which 20 µl was aspirated at 14 µl min^−1^ on a Novocyte flow cytometer (Bucher Biotec, ACEA Biosciences Inc.). Events were collected above thresholds of FSC-H > 150 and SSC-H > 150, and we manually defined gates to distinguish cells from noise based on their green fluorescent signal (see gating strategy in Supplementary figure 5). The absolute cell counts provided by the Novocyte instrument were then converted to cells per mL, taking the intermediate sample dilutions and the fraction of empty droplets measured by microscopy in the emulsion into consideration:

Flow cytometry cell counts after 7 days were taken as the biomass productivity on the respective substrates. We also estimated the maximum biomass of the individual SynCom strains, using their growth in the isolated condition (I-state), by multiplying their relative abundance in the community composition with the total community cell count (Supplementary Fig. 6, see Null-interaction community growth model section below for details).

### Community composition analysis

Community compositions were inferred from sequencing of the amplified V3-V4 region of the 16S rRNA genes in total community DNA, corrected for their known copy number in the individual SynCom strains. Total DNA was extracted from the frozen 10-µl droplet and liquid samples using a protocol adapted from Bramucci et al. (2022)^32^, developed for extraction on small volumes. Briefly, 10 µl of sample was mixed with 7.5 µl of a high pH DTT-lysis buffer (700 µL KOH [0.215 g/10 mL] + 430 µL dithiothreitol [0.2 g/25 ml] + 520 µL of UV-treated (1 h) MilliQ water; resulting pH =12-13) and incubated for 10 minutes (high pH can damage DNA on prolonged incubations) at room temperature before being frozen at -70°C for at least 10 minutes (maximum 4 h). Frozen samples were then fast-thawed at 55°C for 1-2 minutes. 7.5 µl of acid Tris-HCl buffer (‘STOP buffer’, 2.55 M, pH = 5) was then added to neutralise the lysis buffer. Samples were processed in groups of 10 for the thawing and pH-neutralisation steps, to avoid prolonged exposure to the lysis buffer while adding the STOP buffer. After neutralisation, 50 µl of CleanNGS magnetic beads suspension (Clean NA) was added to each lysate to capture the DNA. Beads with bound DNA were washed twice with 200 µl of 80% ethanol on a magnetic rack, after which the tubes were opened for an hour to evaporate the remaining ethanol. Tubes with beads were stored at –18°C overnight until resuspending them in 10 µl of 10 mM Tris-HCl (pH 8.0) to elute the DNA. Eluted DNA samples were transferred to fresh tubes, which were stored at –18°C.

Sequencing libraries were prepared with the Zymo Research Quick-16S Plus NGS Library Prep Kit (V3-V4). This kit uses a one-step PCR protocol with pre-indexed primers (341F and 806R) to amplify the V3–V4 region of the 16S rRNA genes and to simultaneously incorporate Unique Dual Index (UDI) barcodes with Illumina sequencing adapters to the PCR amplicons. The PCR products are pooled in equal volume, and the assembled library is subsequently purified using the magnetic beads provided in the kit. The purified library was sequenced with an AVITI Element Biosciences sequencer at the Lausanne Genomic Technologies Facility, generating up to 100 million 300-bp paired-end reads.

### Sequence read processing and taxonomic identification

Reads were demultiplexed and saved into individual sample FASTA files. FASTA files were processed with CUTADAPT (version 4.4) to remove primer sequences (Forward 1: CCTACGGGDGGCWGCAG; Forward 2: CCTAYGGGGYGCWGCAG and Reverse: GACTACNVGGGTMTCTAATCC), before being handled with the DADA2 (version 1.32.0) R-package for quality control, trimming (parameters used: truncLen = c(260,250), maxN = 0, maxEE = c(4,7), truncQ = 2), merging of forward and reverse reads, generation of the unique sequence variants (ASV) and taxonomic annotation (Silvia database nr99 v138.1 with species training set). We filtered out ASVs that appeared in fewer than four samples in the whole dataset and that encompassed fewer than 100 reads across all samples. We then *grepped* each remaining ASV to a list of reference identifier sequences (from the V3-V4 16S rRNA gene regions from the genomes of the individual SynCom strains) to match with each SynCom member, as described previously^27^. A minor fraction of the sequence reads (3.17 %) could not be attributed and was removed from further analysis. Community compositions were visualised on R (version 4.4.1) with the *ggplot2* package (version 3.5.1). We used *vegan* (version 2.7-1) to compute alpha diversity indexes (Pielou and Shannon), as well as Bray-Curtis distances between samples visualised on NMDS plots.

### Null-interaction community growth model

To compare actual SynCom growth with growth expected from substrate competition alone among the individual strains in the different fragmentation states, we adapted a previously developed mathematical framework that models Monod-kinetic carbon-limited growth from individual species’ growth kinetics (*i.e.*, lag times, maximum specific growth rates, half-maximum constant K_S,_ and yields)^28^. Maximum cell numbers attained for each taxon in the I-state (isolated growth fragmentation condition) were transformed into biomass concentration (fg C mL^−1^) by using a conversion factor of 0.264 pg C for an average *E. coli* cell volume of 2 µm^3^ ^33^ and a volumetric correction based on microscopy images (Supplementary table 1). The strain yield on the used carbon substrate (10 mM succinate = 0.4808 mg_carbon_/mL) was then calculated as the ratio of the formed cell carbon biomass concentration divided by the total starting substrate concentration. The yields thus obtained were above 1 for two strains (Supplementary table 1), suggesting either an overestimation (due to PCR amplification and DNA extraction biases distorting the apparent community composition in the I-state) or that they use another source of carbon substrate. We kept these aberrant yields, with the idea that they would encompass the consumption of any other carbon source beyond succinate that we wouldn’t have considered in the simulation, and would better reflect the community composition observed in the isolated (I-state) fragmented culture condition. Growth rates were inferred as the slope of the linear phase of the ln-OD_600_ growth curves of monocultures (Supplementary Fig. 7), and lag times were deduced from fitting a Monod growth model on these same curves. The model further simulates the starting distribution of cells and species in *n =* 4 000 droplets from random Poisson-sampling (*i.e.*, the expected distribution for droplet-encapsulated cells^31^) of the taxa in the starting mixed inoculum, using droplet volumes matching microscopy observation (Supplementary Fig. 8). The relative strain abundances in the starting inoculum were calculated from the measured flow-cytometry counted relationship of cell densities at their OD_600_ = 1 (which is equivalent to the culture turbidity of each of the individual strains before mixing into the inoculum; Supplementary Fig. 1). Simulated growth for each of the individual droplet SynCom communities is updated for every time step based on the calculation of the utilization of the primary carbon substrate, and is allowed to continue until the available substrate within the droplet volume confinement is too low to sustain further biomass growth. For simulation of bulk liquid culture growth, a single 1-µl volume with 100 000 founder cells (equivalent to the 10^8^ cells/mL in the inoculant, see SynCom assembly and culturing section above) was used with all species in their measured starting proportions.

## Results

### Habitat fragmentation and variations in local starting species richness affect community growth outcomes

Total SynCom sizes measured by flow cytometry of nucleic-acid-stained cells increased 20-100 fold depending on fragmentation (droplet emulsion or bulk) and media conditions (Fig. 1b, succinate or SE). Growth on succinate was slightly slower than on SE, but community sizes attained after 7 days in numbers of cells per unit of (collapsed) volume were similar, suggesting similar carrying capacities between the two media (Fig. 1b). The starting community size measured in the collapsed droplet emulsions at Day 0 was systematically lower than calculated from diluted, mixed and quantified individual strains in the inoculum, suggesting that cell densities measured from broken emulsions are potentially underestimated (Fig. 1b). Community sizes were different among fragmentation states, the highest being attained in low-order growth on both succinate (1.3×10^9^ cells/ml) and soil extract (9.8×10^8^ cells/ml, p < 0.05 versus all conditions, HSD Tukey test). This suggests a positive effect of habitat fragmentation on bacterial community growth, regardless of the nutrient context and when normalizing the community sizes to the actual proportion of droplets exhibiting growth (*i.e.*, excluding culture volume of empty droplets; Fig. 1c and Supplementary Fig. 3). In comparison, growth in the bulk liquid suspension on SE was lower than in the droplet conditions, but was higher on succinate (Fig. 1c). Succinate samples showed high replicate variability which might have been due to the formation of cell aggregates during (standing) incubation. Repetition of the F-condition with independently prepared SynCom inoculum on both media with more vigorous sample resuspension indeed lowered the replicate variability (Fig. 1c, condition F2), attaining community sizes equivalent to the H-droplet condition. Of note, however, that bulk community counting is not completely comparable with the procedure of counting cells in broken droplet emulsions, because of the absence and removal of the PFO-oil.

Cell growth measurements in individual droplets, approximated by the total contrast change score (CCS, Supplementary Fig. 3) indicated rapid growth in part of the droplets particularly for the succinate condition, followed by a gradual increase for the remainder of the droplets (Fig. 1d). The proportion of droplets exhibiting growth increased over time in all conditions (Supplementary Fig. 3), as expected both from the Poisson-random cell distribution at start (in the I-case, with most droplets being empty) and variable growth depending on the per-droplet species starting mixture. For example, 5, 40 and 75 % of droplets showed detectable cell biomass (CCS above the 95^th^ percentile of medium droplets alone) after one day in I-, L- and H-conditions on succinate, respectively, which increased to 22, 95 and 96 % after 7 days (Supplementary Fig. 3c). Cell biomass in SE-droplets increased from 12, 62 and 80 % to 40, 87 and 99 %, for I-, L- and H-order conditions, respectively, between Day 1 and 7 (Supplementary Fig. 3d). Under I-conditions, a larger number of empty droplets is present from the start, which contributes to the generally lower fraction of droplets with detectable growth (Supplementary Fig. 2 & 3).

For succinate, the heterogeneity in droplet productivity (detected as the variation in the CCS score, Fig. 1e) kept increasing over time in case of isolated growth, while it leveled off after one day in low- and high-order growth conditions, followed by a slight decrease (Fig. 1e). Growth in SE droplets was more gradual than on succinate (Fig. 1d) and the CCS heterogeneity among droplets increased over time (Fig. 1e). A higher CCS heterogeneity is indicative for a wider variation in growth among individual droplets, in agreement with our hypothesis that randomly divided sub-communities would show more diverse growth in SE medium (where multiple nutritional niches are available) than on succinate. The high CCS heterogeneity in isolated growth conditions is expected from the differences in the ability of the individual species to grow on the carbon substrates (Supplementary Fig. 7).

Altogether, these results showed how different degrees of habitat fragmentation and the resulting variations in species and cell numbers at the start impact the growth dynamics and sizes of the attained communities, both at the level of individual droplets and the community as a whole. For both substrate conditions in droplet emulsions, intermediate fragmentation (L-condition) showed the highest aggregate community productivity.

### Varying habitat fragmentation and local species order lead to different global community compositions

To understand whether the observed growth differences of the SynComs incubated under the varying fragmentation regimes would also result in compositional changes, we analysed individual population growth over time from species-specific 16S rRNA gene amplicon variants (relative abundances) and total community flow cytometry cell counts.

Relative abundances after 7 days indicated clear and consistent compositional differences both as a function of medium type and fragmentation regime (Fig. 2a). Community compositions at day 7 showed a gradual change between the two extremes of the I- (1–2 cells/taxa per droplet) and the F-state (full community in a homogenous mixture). Community compositions across all samples indeed diverged after day 1, depending on the fragmentation regime, on both media (*i.e.*, I, L, H or F; permanova – effect of Condition, R^2^ = 0.28 and R^2^ = 0.31 for succinate and SE, respectively, p = 1×10^-4^, Fig. 2b). Species relative abundances in the merged communities (*i.e.*, broken droplet emulsions) over time visually changed as an effect of fragmentation state (Supplementary Fig. 9). For example, succinate-grown communities after one day became dominated by both *Pseudomonas* strains, except in the I-case of 1–2 cells and species per droplet (Supplementary Fig. 9). *Flavobacterium* dominated early growth on SE in the F-condition, whereas it was the major fraction in the I-condition at the end (Supplementary Fig. 9). At day 7, some of the more abundant strains in the I-state (*e.g.*, *Mesorhizobium* on succinate, or *Flavobacterium* on SE) decreased in proportion along the fragmentation gradient from the L- to the H- and the F-state, whereas *e.g.*, *Lysobacter* in SE increased in proportion from the I- to the L- and the H-state before decreasing again in the F-state (Fig. 2a).

**Figure 2:**
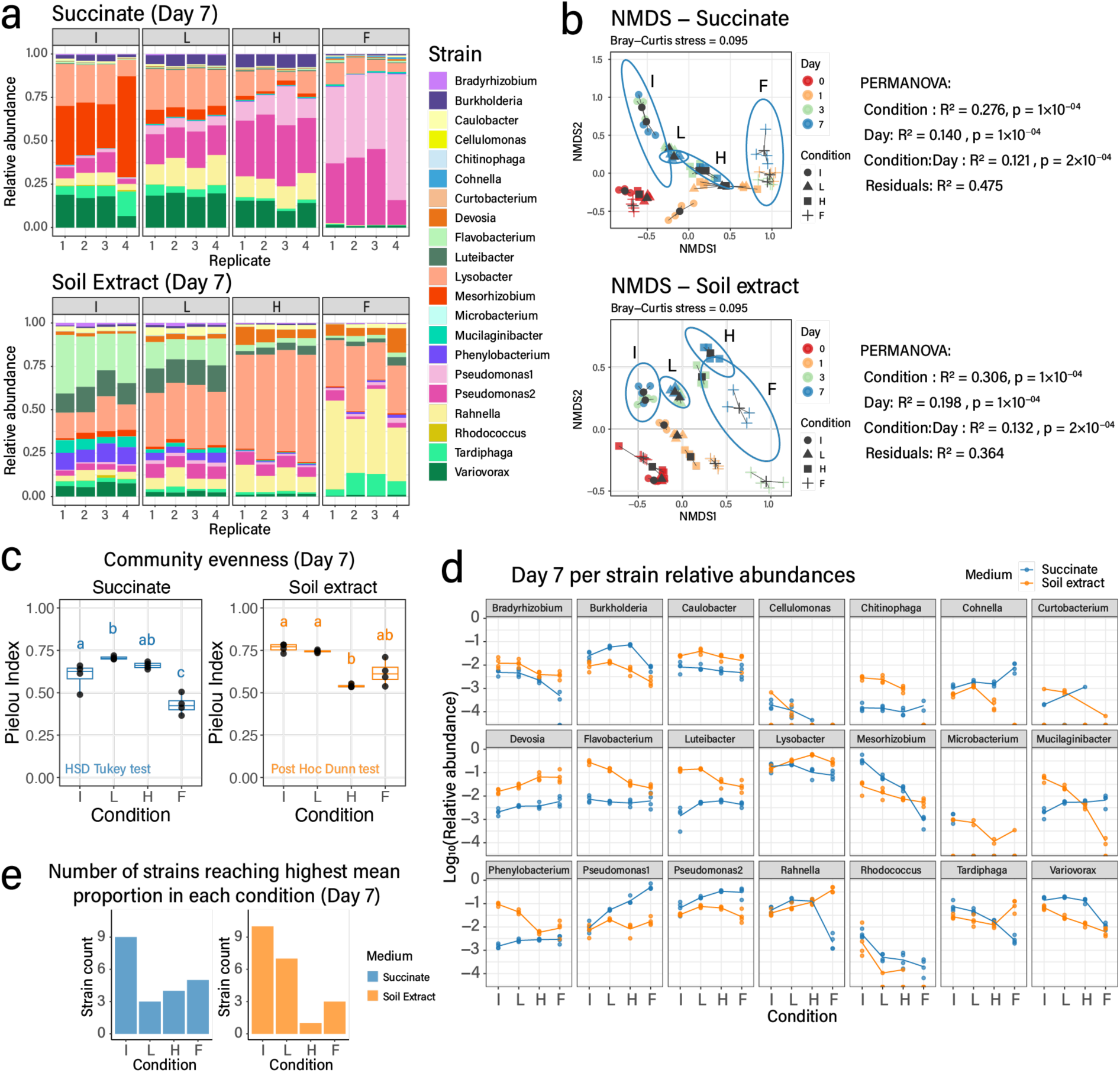
Community composition changes under different habitat fragmentation. **a** SynCom composition after 7 days on succinate (top) or SE (bottom) for each of the fragmentation conditions. Stacked bar plots show relative strain abundances (colors according to the legend on the right) for each of the replicates per condition. **b** Clustering of SynCom compositional trajectories by habitat fragmentation on succinate and on soil extract (SE). Plots show non-metric multidimensional scaling (NMDS) of Bray-Curtis dissimilarities of SynCom strain relative abundances determined by 16S rRNA gene amplicon sequencing. Colors and symbols depict time points and fragmentation conditions as per the legend. Ellipses group fragmentation conditions at Day 7. I, L, H, and F as in Fig. 1A. **c** SynCom evenness measures (Pielou index) at Day 7 per fragmentation conditions and growth substrate. Letters group statistically indifferent values from Anova followed by post-hoc Tukey test (succinate, p-value < 0.05) or by post-hoc Dunn test (soil extract, p-value < 0.025). Boxplots indicate the lower, upper quartiles, and median. Black dots show individual replicate values. **d** Trends for individual strain relative abundances as a function of decreasing fragmentation state. Lines connect the mean log_10_-transformed relative abundances at Day 7 with circles indicating individual replicate values, and colors, the growth substrate. **e** Strain mean optimal relative abundance per growth substrate and fragmentation condition. Bars show the number of strains where their mean relative abundance (from four replicates) is the highest.

Both the community evenness (Fig. 2c, Pielou index) and diversity (Supplementary Fig. 10, Shannon index) indices at day 7 were highest for the L- and H-conditions on succinate, and the I- and L-conditions on soil extract. They were significantly lower for the F-conditions on succinate, and both H- and F-conditions on soil extract (p < 0.05 for HSD Tukey test, and p < 0.025 for Post Hoc Dunn test). This would imply that low-order taxa combinations in fragmentated habitats are positive for global-scale community taxa coexistence, in particular on substrate complex conditions. In contrast, as mentioned above, SynComs in the I-condition tend to have a lower productivity than the L-state, which, at least for the soil extract, is statistically significant (Fig. 1c, p < 0.05, HSD Tukey test). The lower SynCom diversity in the H- and F-conditions may be the result of an increasing impact of competition, as reflected by the dominance of *Lysobacter*, *Rahnella,* and *Pseudomonas* (Fig. 2a & 2c).

More in general, individual relative strain abundances either followed a constant ‘downward’ trend from I- to F-conditions, a constant upward, or a bell-shaped trend (Fig. 2d), suggesting that they react in varying ways to increasing species orders and decreasing fragmentation. The I-condition was optimal for the highest number of species in both media (nine species on succinate and ten on SE, Fig. 2e). The second most preferred regimes were the F-condition on succinate and the L-condition on SE, suggesting that fragmentation might be advantageous to a larger number of species when the number of nutritional niches increases (Fig. 2e). This could be the consequence of several strains growing poorly by themselves on succinate but reaching higher proportions in the full-community mixture where they may benefit from leaked metabolites from dominant members (*e.g.*, *Cohnella*, *Devosia*, *Mucilaginibacter*, Supplementary Fig. 7, Fig. 2a and 2d).

These results thus demonstrated how different degrees of habitat fragmentation and associated species orders affect compositional outcomes at the global community level. Notably, lower starting species numbers in small volume habitats tend to favor the coexistence of species at the community level, which otherwise would be outcompeted.

### Faster-growing species dominate communities with decreasing fragmentation on single but not on mixed carbon substrates

Next, we tried to understand the underlying causes for the differences in taxa growth as a function of fragmentation and substrate condition. We first quantified the effect of the fraction of the total accessible habitat for each species. The total species-accessible habitat depends on the fragmentation condition, because individual droplets do not contain all taxa (Supplementary Fig. 2), and each strain therefore has access to only a fraction of the total habitat medium volume and substrate. This contrasts with the F-condition, where all strains in the mixture at the start have access to the complete medium volume and its substrate. Any observed differences in relative abundances (particularly for strongly competitive strains in the community) may thus simply be a direct reflection of the strains having access to different fractions of the total habitat, and not necessarily of them changing their competitive behavior. To follow this idea, we estimated the theoretical starting species proportions and their accessible habitat fraction under the different fragmented conditions (Supplementary Fig. 2f). Then we plotted the final observed relative abundances for each of the strains as a function of the accessible habitat fraction (Fig. 3a). Since this is expressed as a continuum of accessible habitat, the trends for individual strains and substrate conditions can be formalized by regression lines (Fig. 3a).

**Figure 3:**
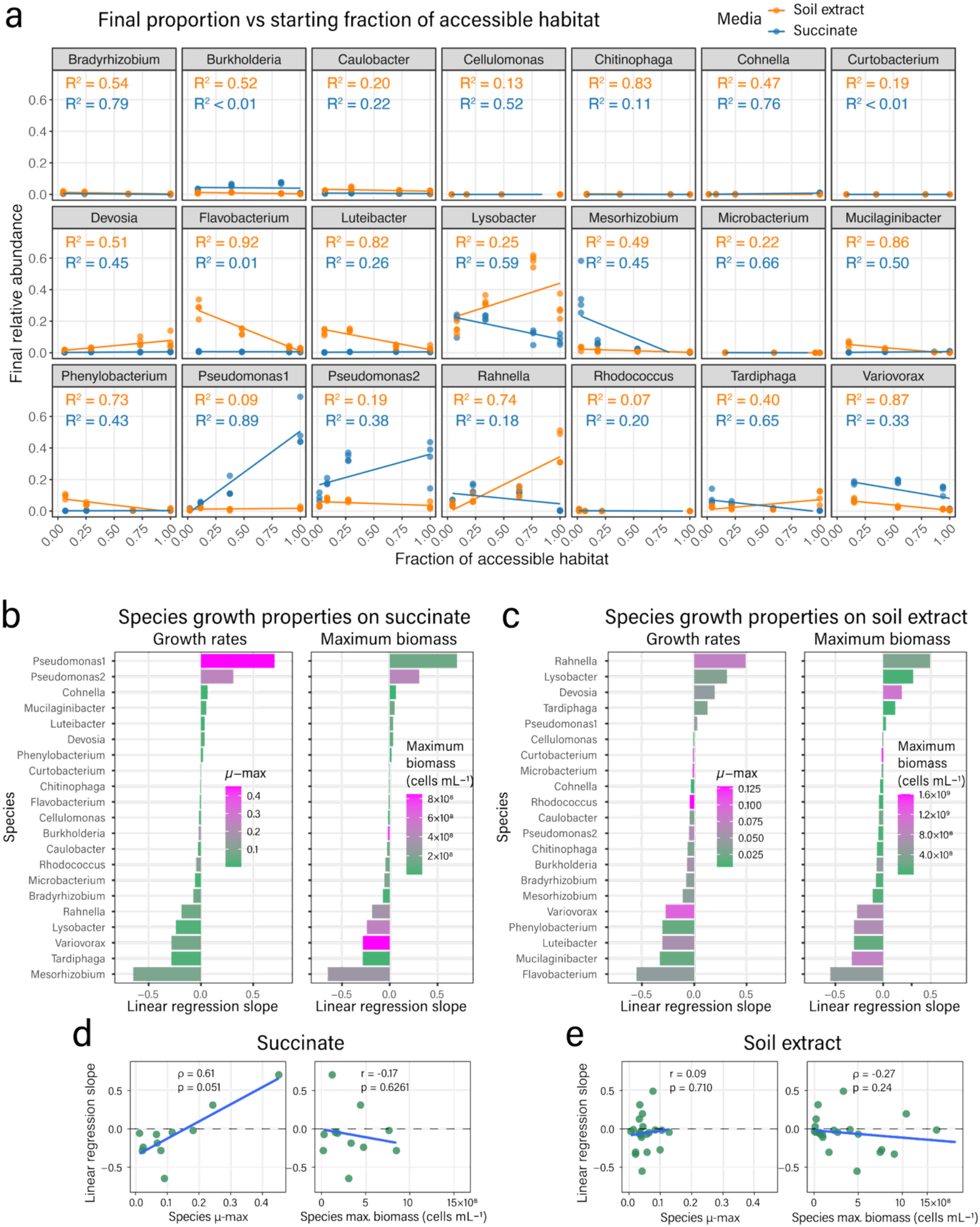
Relation between growth kinetics of individual SynCom strains and their response to decreasing habitat fragmentation. **a** Square-root-transformed final relative abundances of each SynCom member as a function of the square-root-transformed fraction of accessible habitat under each of the fragmentation conditions (*i.e.*, from I – isolated, to L, H and F, full mixture; F having a fraction of 1). Lines denote linear regression with corresponding R^2^ values displayed in each facet. Orange, values on soil extract; blue, succinate. Circles show individual replicate values. Ranked linear regression slopes from (a) for succinate (**b**) or soil extract (**c**), overlaid by a color scale indicating the mean monoculture growth rates or their maximum cell numbers measured in bulk (*n* = 3 replicates). Correlations between individual maximum specific growth rates (µ_max_) or maximum cell numbers in bulk and their responses to decreasing habitat fragmentation (*i.e.*, the slopes from (a)) for (**d**) succinate or (**e**) soil extract. r and ρ values are respectively Pearson and Spearman correlation coefficients with associated p-values from correlation statistical testing. To test correlation, only strains for which a net growth was detectable after 7 days in liquid monoculture were considered (see Supplementary Fig. 7)

For six (succinate) and five (SE) out of the 21 strains, an increasing accessible habitat fraction resulted in increased relative abundances (Fig. 3a). Only two of those strains showed increasing trends under both media conditions (*Devosia* and *Pseudomonas* 1). Around one-third of the strains were more or less indifferent to increasing accessible habitat fraction and maintained a constant fraction in the community (linear regression slopes close to 0, Fig. 3a-c). Some of those (*e.g., Cellulomonas*, *Chitinophaga,* or *Curtobacterium*) grew rather poorly on both media, whereas others (*e.g.*, *Burkholderia* and *Caulobacter*) maintained relative abundances of ca. 1-5% under all conditions (Fig. 3a and 2d). Finally, several strains decreased in relative abundance upon increasing accessible habitat volume (*e.g.*, *Mesorhizobium* and *Tardiphaga* on succinate, or *Flavobacterium* and *Mucilaginibacter* on SE, Fig. 3a).

The most competitive strains, judged from the slope of the relative abundance versus increasing accessible habitat fraction, were *Pseudomonas* 1 and *Pseudomonas* 2 on succinate, and *Rahnella* and *Lysobacter* on SE. The poorest competitors (with the highest negative slopes) were *Flavobacterium* on SE and *Mesorhizobium* on succinate, which also indicates that these strains grow well by themselves but poor in competition with others (Fig. 3a). The high degree of linearity and positive slope in the some of the regressions (*e.g.*, *Pseudomonas* 1 on succinate, R^2^ = 0.89, Fig. 3a), suggests that these strains are essentially limited by accessibility to the total habitat volume, whereas high linearity with negative slope (*e.g.*, *Flavobacterium* on SE, R^2^ = 0.92, Fig. 3a) is indicative for the hindrance by increasing number of species. The linear fit slopes can thus be used as an indicator to rank the species’ competitiveness (Fig. 3b and 3c) and can be compared to the determined growth rates and maximum observed cell numbers of each strain in isolation in liquid suspension (Supplementary Fig. 7). This showed for succinate a positive correlation of slope to growth rate yet borderline significant (Spearman coefficient ρ = 0.61, p = 0.051, Fig. 3d), with the fastest growing species (*Pseudomonas* 1 and *Pseudomonas* 2) being the top ranked in slope, but not to maximum observed cell numbers (Pearson coefficient r = -0.17, p = 0.626, Fig. 3d). On SE, in contrast, neither growth rate nor maximum observed cell numbers could explain the observed relations between accessible habitat fraction and relative abundance (Fig. 3c and 3e).

While the regression values of the linearly fitted trendlines were high for some strains, they were relatively poor for many others (*e.g., Pseudomonas* 2 and *Rahnella* on succinate or *Lysobacter* and *Tardiphaga* on SE, Fig. 3a), echoing the above-mentioned ‘bell-shape’ in the relative abundance variation along the increasing species order (from I to F, Fig. 2d). This nonlinearity suggests that additional interaction effects, such as metabolic cross-feeding, influence the relative abundances of those species in the mixtures. This nonlinearity could also stem from a trade-off between a beneficial spatial isolation of competitors, but a limited access to the whole volume and resources for the intermediate fragmentation degrees (*i.e.*, L- or H-states).

### Spatial partitioning and substrate competition are the main drivers of community assembly on single carbon substrate

To better understand how much spatial fragmentation alone is contributing to the variation in community composition, we simulated community growth in the absence of interspecific interactions. We assumed here that community growth is the sum of Monod growth of the constituting individual populations, constrained by the limiting amount of total available substrate and habitat volume, and the starting taxa proportions in individual droplets. We only focused on the case of succinate because its measured monoculture growth kinetics better reflect the single substrate utilisation than in the case of SE, which contains a mixture of different substrates. Maximum growth rates and lag times were fitted from the observed increase in monoculture turbidity in 96-well plate measurements of succinate growth (Supplementary Fig. 7). Strain yields were inferred from their absolute abundances under the I-condition (strains in isolation; Fig. 2a, relative abundances from sequencing multiplied by the total cell counts in the merged droplet conditions; Fig. 1c, Supplementary Fig. 6). The number of starting cells for each strain (see starting proportion in Supplementary Fig. 1) was simulated by Poisson-random subsampling of the community in droplets of either 43 µm or 67 µm diameter, assuming similar starting cell densities as estimated from imaged SYBR-stained inoculum encapsulated in droplets (1.74×10^8^ cells/mL for the L-, H- and F-, and 1.74×10^7^ cells/mL for the I condition, Supplementary Fig. 2).

For the isolated (I-case) fragmentation condition, the predicted and observed relative abundances in the SynCom at day 7 were very similar, as expected (‘resource×1’, Fig. 4a), indicating that the community growth on succinate, in that case, is indeed a sum of the growth of the individual populations. In contrast, the simulated composition under the higher species order conditions (L, H and F cases) diverged drastically from the observed relative abundances (‘resource×1’, Fig. 4a). Indeed, the fold increase of community size was highly underestimated (1.1-to-7.5-fold instead of 10- to 100-fold observed increase, Fig. 4b and 4c), leading to very little change from the starting composition (Supplementary Fig. 11). These differences may point either to an underestimation of the species yield on succinate, or to the effect of interspecific interactions (such as cross-feeding), which were not accounted for in the simulations^28^.

**Figure 4:**
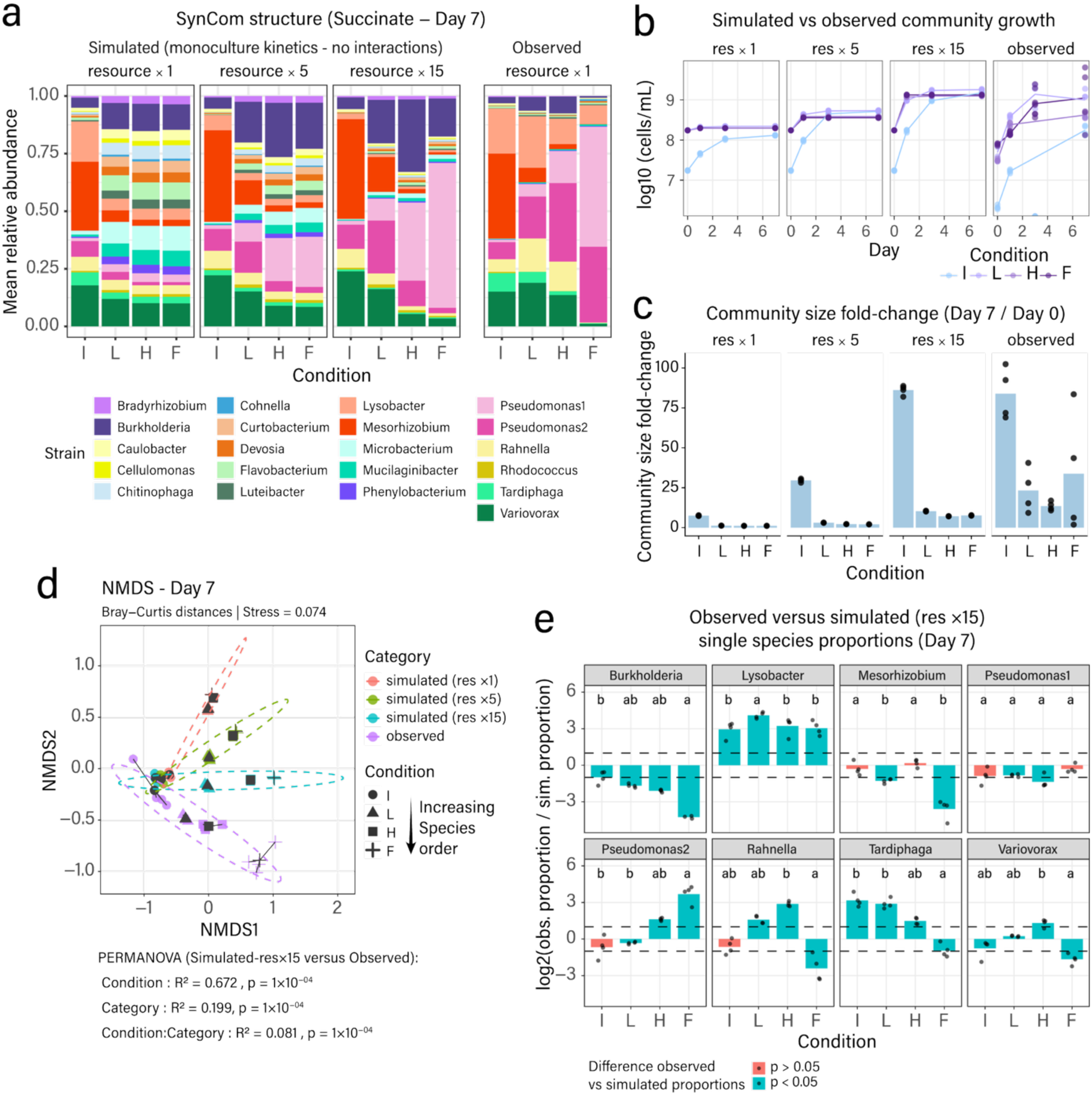
Simulation of SynCom growth on succinate as the sum of individual population growth without interactions. **a** Observed and simulated mean relative species abundances after 7 days of growth under different fragmentation (I, L, H) or bulk (F) conditions. Simulations use a Monod substrate utilization growth model for each of the individual SynCom strains and their growth kinetic parameters (Materials and Methods). Stack bars show the mean relative abundances from *n* = 4 independent replicates, derived from 4000 individual simulated droplets or 1 bulk culture (F) each. Resource ×1, simulations with original succinate concentration; Resource ×5, or ×15, simulations with five or fifteen times higher succinate concentration to allow more generations of growth. **b** Simulated versus observed SynCom growth kinetics. **c** Simulated versus community size fold-change between day 0 and day 7. **d** Non-metric multidimensional scaling representation of the simulated and observed community composition dissimilarities computed from Bray-Curtis distances. Replicate community dots are connected to a geometric centroid in dark grey. Shapes indicate the different fragmented states, and colours indicate simulations or experimental observations. Ellipses combine different conditions at a 95 % confidence interval. Permanova analysis shows the contribution of the fragmented state (Condition) or interspecific interactions (Category, which accounts for the difference between observed and simulated community compositions), using the observed and simulated Resource×15 community composition. **d** Log_2_-transformed mean ratio (bars) of observed and simulated relative species proportion at day 7 (n = 4 simulations and experimental replicates, see Supplementary Fig. S12). Bars in blue point to significantly different ratios (p < 0.05, two-sided t-test or Wilcoxon rank sum test), with letters indicating significance groups in Anova followed by HSD Tukey post-hoc testing (p < 0.05) or Dunn testing (p < 0.025). Dashed lines indicate the two-fold difference threshold. Only species reaching a minimum observed proportion of 5% of the community in at least one of the fragmented states are shown.

We repeated simulations with five- and fifteen-fold succinate concentration increase without changing starting cell numbers to allow a simulated number of generations closer to our experiment in the I-condition (Fig. 4b and 4c). In this case, the simulated compositions more closely resembled the experimental observations (Fig. 4a, 4b and 4c), recapitulating some of the observed strain-dominance patterns such as by *Pseudomonas* 1 from I- to F-condition (Fig. 4a). PERMANOVA analysis on Bray-Curtis dissimilarities between the observed and simulated communities with 15-fold increased succinate concentration indicated that the state of fragmentation explained 67 % of the variation in community structure (‘Condition’, p-value = 1×10^-4^, Fig. 4d), whereas Category (observed or simulated) explained only 20% of variation (p-value = 1×10^-4^, Fig. 4d). These results suggest that differences in spatial segregation of species (*i.e.*, seeding effects) and emerging substrate competition are the main drivers for alternative community outcomes under variable fragmentation conditions.

Looking more closely at simulated individual species proportions in the last simulation (‘resource × 15’), we wondered whether specific fragmentation states would favor the emergence of (positive) interspecific interactions. To understand this, we inspected how the ratio between observed and simulated proportions varied across fragmentation conditions for the most abundant species (*i.e.*, with an observed proportion > 5% in any condition, Fig. 4e). Several clear differences indeed remained between simulated and observed relative abundances (Fig. 4a, 4e, and Supplementary Fig. 12), which could not be explained by individual growth kinetic differences on succinate and may be attributable to interspecific interactions. For some of those, observed and simulated relative abundances as a function of accessible habitat varied in the same way, whereas for others, simulated trends differed from observed ones, for example, *Rahnella* and *Pseudomonas* 2 (Supplementary Fig. 12). The ratio of observed and simulated relative abundances, which can translate into the direction and strength of interspecific interactions, were statistically significantly different for 6 out of 8 species, at least between the three mixed-species conditions (L, H and F, Fig. 4e). This suggested that the degree of fragmentation can modulate the unfolding of interactions at the global community level in a species-dependent way (Fig. 4e).

## Discussion

Our understanding of the processes that drive and control bacterial community development is still limited. In particular, the role(s) of the habitat spatial structure and the degree of habitat fragmentation have not been studied in depth, which may be due to the technical difficulty of producing controlled habitat fragmentation conditions. We hypothesized that fragmentation should create a bottleneck on the number of co-occurring taxa when the dimension of micro-habitats is small enough, which we expected would affect global community development. To test this experimentally, we took advantage of picolitre droplet culturing systems. By using a 21-membered synthetic community composed of culturable soil isolates, we found that composition outcomes change gradually as a function of habitat fragmentation and the imposed local starting species order, enabling different strains to thrive under particular fragmentation states. The relative abundance of some strains in the community was directly and linearly dependent on the fraction of fragmented habitat they had access to. We consider such strains as being strong opportunistic competitors that manage to increasingly dominate as they gain access to more habitat, independently of increases in the number of co-inhabiting taxa. For others, an increasing number of local neighboring taxa resulted in a decline in their relative abundance, indicating a loss of competitiveness. Despite this, community growth simulations based only on individual monoculture kinetic parameters indicated lower than observed overall community size fold-change, suggesting that, on average, individual population growth in a community is favored by utilization of additional substrates generated because of metabolism of primary substrates. Simulations further confirmed growth competitive trends for some species and additional emerging interaction effects for others, and a general dependency of community composition on habitat fragmentation.

We expected that increasing habitat fragmentation would gradually isolate taxa and restrict cells from interacting in the community, limiting the importance of emerging interactions at higher species orders. A pioneering study in this field using communities of engineered *E. coli* strains showed that increasing fragmentation lowered global diversity in the case of strains being obligatory cross-feeding, but increased diversity if strains in the community were characterized by resource competition^23^. In our studies, we observed the highest alpha diversity for the highest degrees of fragmentation (I- and L-conditions), where the average per-droplet number of species is low (Fig. 2). This would indicate that habitat fragmentation and low taxa orders facilitate species coexistence and favor global (*i.e.*, that of the aggregate community among the complete habitat) community evenness. In contrast, larger-volume (bulk) habitats and high taxa orders tended to result in outgrowth of fast-growing strains, and skewed global community evenness^34^. This evenness effect occurred both for succinate and for SE as growth media, indicating that SynCom growth is dominated by substrate competition. Simulated community growth from the sum of individual growth on a single primary substrate (succinate) in all fragmented conditions indeed led to less generations than experimentally observed, suggesting that utilization of metabolic byproducts is an essential element of community development, irrespective of competitive interactions (Fig. 1 and Fig. 4)^18^. Including cross-feeding in growth models has previously better explained co-culture dynamics of competing strains^28^, but is difficult to implement on higher-order species communities in the absence of interaction parameters.

On closer inspection, our results suggest that individual SynCom species are differently adapted to varying degrees of habitat fragmentation. Each of the imposed fragmentation states and resulting starting species distributions was optimal for at least one SynCom member (Fig. 2). This suggests that different taxa have evolved to operate optimally under different ranges of micro-habitat dimensions and local species-order. For example, fast-growing species take proportionally more advantage of large habitats (*e.g., Pseudomonas* 1), whereas slower-growing species more optimally prevail in small, insulated micro-habitats (*e.g., Mesorhizobium*). If different degrees of fragmentation apply such an evolutionary pressure for different traits on microbial taxa, one could question in the future perspective whether ecological strategies in nature indeed spatially correlate with a certain range of spatial habitat dimensions. This could also imply that a continuum of ecological strategies^35^, connecting the life-history archetypes covered by the classical r-K (fast growth – slow growth and adaptability) or YAS frameworks (high yield – resource acquisition – stress tolerance)^36^, is enforced under a continuum of habitat dimensions, which influence both the local taxa order and impose spatial and substrate restrictions.

Our observations also place a different perspective on the view that high-order interactions have a positive effect on species coexistence, and have often been cited as important contributors to the natural taxa diversity^37–39^. The role of high-order interactions, however, is difficult to assess and has often been inferred solely from fitting empirical community compositional variations in modelling frameworks that included higher-order terms in their equations^38–41^. To experimentally quantify the role of higher order interactions requires decomposing communities into smaller order combinations (*e.g.*, alone, pairs, triplets, etc)^2,5^, culturing these under the imposed conditions, and observing resulting community structures. Unfortunately, the number of combinations increases factorially with the number of starting strains, and is physically unrealistic for consortia with more than 20 strains. Aguadé-Gorgorio and Kéfi recently demonstrated that bacteria can coexist even in the absence of higher-order interactions, suggesting their role may have been overestimated^42^.

By fragmenting a 21-membered community to different degrees and parallel culturing of droplet-incubations, we directly observed the effects of species starting order both on community composition and productivity. Can we conclude that variable local species orders are enabling different emergent interaction effects (higher-order interactions) that alter the community composition at the global scale? Or should we conclude that the observed differences in community compositions are the result of the imposed fragmentation conditions and the starting compositions (without the necessity to invoke emerging interactions)? To answer this question, we compared our empirical data with community growth simulations, which exclude emerging interspecific interactions. The simulations produce the starting species conditions and their relative abundances through Poisson-random sampling into small culture volumes, and then simulate individual population growth under carbon limitation from empirically measured individual growth kinetics on the same substrate. According to this comparison, the proportion of variation explained solely by inherent growth kinetic differences among strains on community outcomes is 67% (Fig. 4). Interspecific interactions, in contrast, made only a modest contribution to succinate community growth (R^2^ = 0.20, Fig. 4). We did find evidence for emerging interspecific interactions, for example, in the statistically significant differences of simulated compared to actual relative abundances of *Pseudomonas 2* in the F-condition (Fig. 4), but there was no consistent effect of the imposed species order (*i.e.,* L, H and F) and the direction of the ratio of the simulated and observed relative abundances. This suggests that it is the physical fragmentation itself (or insulation), more than the resulting starting species order, that determines how interactions unfold in the system (Fig. 4). Habitat fragmentation thus directly affects the degree of species intermixing and constrains the extent to which each of them can multiply. Although this creates variation in the emerging interaction patterns in the physically segregated starting communities (Fig. 2 and Fig. 4), it is primarily the chance of finding itself in a pristine colonisable habitat to be shared with others rather than the degree of emerging interactions that controls the dynamics of microbial community growth.

The micro-droplet emulsion approach allowed for recreating micro-fragmented conditions, which represent fragmentation in natural habitats to some extent, while allowing good control over starting conditions. For example, partially water-saturated soil porosity covers the range of micro-habitat dimensions like those tested here (*e.g.*, 10-100 µm diameter size)^12^, but soils typically go through dynamic changes of increased and diminished connectivity between micro-habitats as a consequence of rainfall and drying^43^. Soil microcosms can be used to grow communities under fragmented relevant conditions^27^, but in contrast to micro-droplets, offer little control over the local starting distribution of species and the potential for interactions between micro-pockets to develop (*i.e.*, the degree of connectivity). Future research may expand the concept of dynamic connectivity in droplet emulsion incubations by alternatingly fragmenting and growing, then merging and re-fragmenting communities during multiple cycles. This could allow for investigating meta-community dynamics^44^ and illustrating the effects of microhabitat seeding processes on community outcomes.

We previously showed how microhabitat confinement can lead to alternative interaction outcomes between pairs of species, originating from physiological heterogeneities of founder cells^22^. Here, we highlight a different process with important ecological implications for the assembly of multi-species communities, namely the degree of species order interactions from random starting assemblies within physically constrained micro-environments, primarily limiting fast opportunistic growth and enabling higher richness and evenness to be sustained. This principle may also be used for synthetic community engineering to control strain relative abundances or for the selection of more optimized bacterial inocula for microbiome interventions in natural habitats with their specific fragmentation characteristics.

## Acknowledgments

The authors wish to thank Senka Causevic for her initial help setting up SynCom assembly and cultivation, Prajwal Padmanabha, Alessia Del Panta, and Massimo Amicone for their helpful advice on statistical analyses and simulations to perform, as well as Simon Yersin and Garance Sarton-Lohéac for instructing 16S rRNA gene amplicon sequencing read processing.

## Author contributions

M.B., A.P., and J.M. conceived the studies and designed experiments. M.B. and A.P. performed all droplet and liquid culture experiments, DNA extraction, and 16S rRNA gene amplicon library preparation. M.B. analyzed image, flow-cytometry and sequencing data. I.G. wrote the simulation model. M.B. conducted simulations. M.B. and J.M. wrote the draft manuscript. All authors gave input, verified, and corrected the written manuscript. J.M. acquired funding and coordinated the work.

## Data and code availability

Source data are provided with this paper. All raw imaging data, processed droplet productivity data, raw and processed sequencing fastq files, flow-cytometry processed cell counts data, and numerical source data values underlying figure elements, as well as R/MATLAB/PYTHON scripts and code used for subsequent analyses, are available from a single downloadable link on Zenodo (https://doi.org/10.5281/zenodo.18468666) under accession number 18468666^45^.

