## Supplementary material for "Habitat fragmentation controls bacterial community composition outcomes"

**Supplementary Tables:**

Supplementary Table 1: Single-carbon biomass values per species' cell and yields measured on succinate

**Supplementary Figures**

Supplementary Figure 1: Quantification of the SynCom inoculum composition

Supplementary Figure 2: Projection of the starting species distributions in droplets

Supplementary Figure 3: Droplet image analysis

Supplementary Figure 4: Correlation between resorufin red fluorescent signal and bright field CCS.

Supplementary Figure 5: Gating strategy to identify SynCom cells among flow-cytometry events.

Supplementary Figure 6: Absolute species abundances after 7 days of SynCom growth

Supplementary Figure 7: Monoculture bulk liquid growth kinetics

Supplementary Figure 8: Size distribution of droplets after cell encapsulation at Day 0

Supplementary Figure 9: Compositional variation of the SynCom over time in the different media and conditions

Supplementary Figure 10: SynCom alpha diversity

Supplementary Figure 11: Simulated compositional variation of the SynCom over time on succinate.

Supplementary Figure 12: Observed and simulated species proportions

**Supplementary Table 1: Single-carbon biomass values per species' cell and yields measured on succinate**

| SynCom strain | Dry carbon biomass (fg/cell) | Yield measured in monoculture assay (flow-cytometry) | Yields inferred from fragmented I-state (isolated) |
|---|---|---|---|
| *Cellulomonas* | 250 | 0,0095 | 0,0019 |
| *Bradyrhizobium* | 250 | 0,0162 | 0,0174 |
| *Luteibacter* | 250 | 0,0579 | 0,0055 |
| *Phenylobacterium* | 250 | 0,0031 | 0,0048 |
| *Mesorhizobium* | 250 | 0,1618 | 1,6470 |
| *Lysobacter* | 275 | 0,2737 | 0,5440 |
| *Caulobacter* | 275 | 0,0756 | 0,0273 |
| *Devosia* | 250 | 0,0364 | 0,0056 |
| *Cohnella* | 250 | 0,0022 | 0,0109 |
| *Tardiphaga* | 250 | 0,0119 | 0,3121 |
| *Rahnella* | 275 | 0,1929 | 0,2454 |
| *Rhodococcus* | 250 | 0,0827 | 0,0674 |
| *Burkholderia* | 290 | 0,4615 | 0,0705 |
| *Chitinophaga* | 500 | 0,1156 | 0,0007 |
| *Variovorax* | 275 | 0,4863 | 0,2332 |
| *Microbacterium* | 250 | 0,0998 | 0,0013 |
| *Flavobacterium* | 250 | 0,0250 | 0,0116 |
| *Mucilaginibacter* | 250 | 0,0619 | 0,0039 |
| *Curtobacterium* | 250 | 0,0279 | 0,0005 |
| *Pseudomonas* 1 | 290 | 0,0746 | 0,0956 |
| *Pseudomonas* 2 | 290 | 0,2678 | 1,0789 |

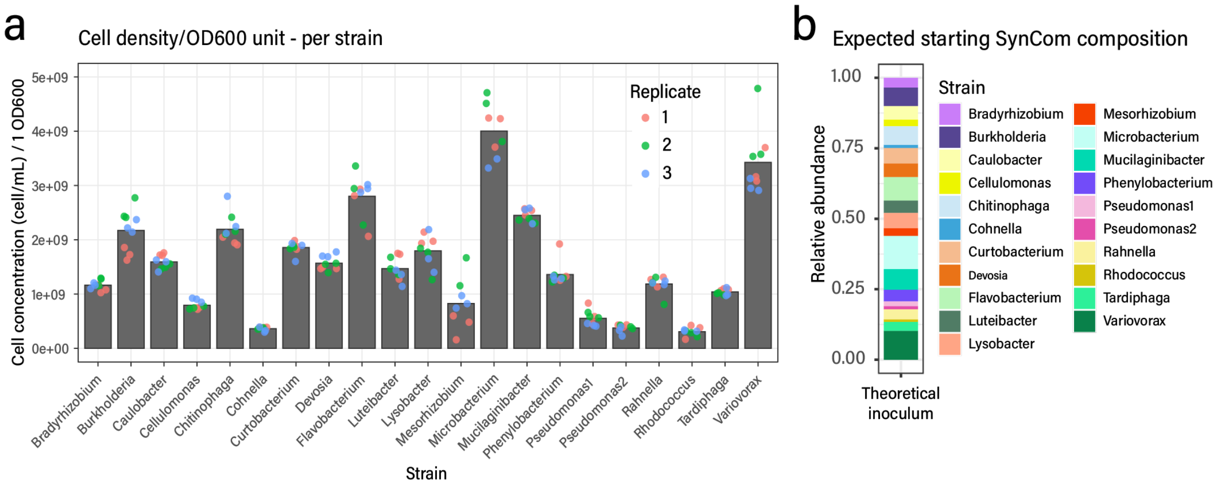


**Supplementary Figure 1: Quantification of the SynCom inoculum composition. a**) Cell densities measured by flow cytometry on triplicate mono-species suspensions at $OD_{600}$ = 1, as used to prepare the SynCom inoculum in all experiments. Individual dots correspond to flow cytometry technical replicate measurements. **b**) SynCom inoculum composition deduced from the cell densities measured in a). The deduced composition was used to predict the fraction of emulsion accessible to each species following encapsulation in droplets (see Supplementary Fig. 2).

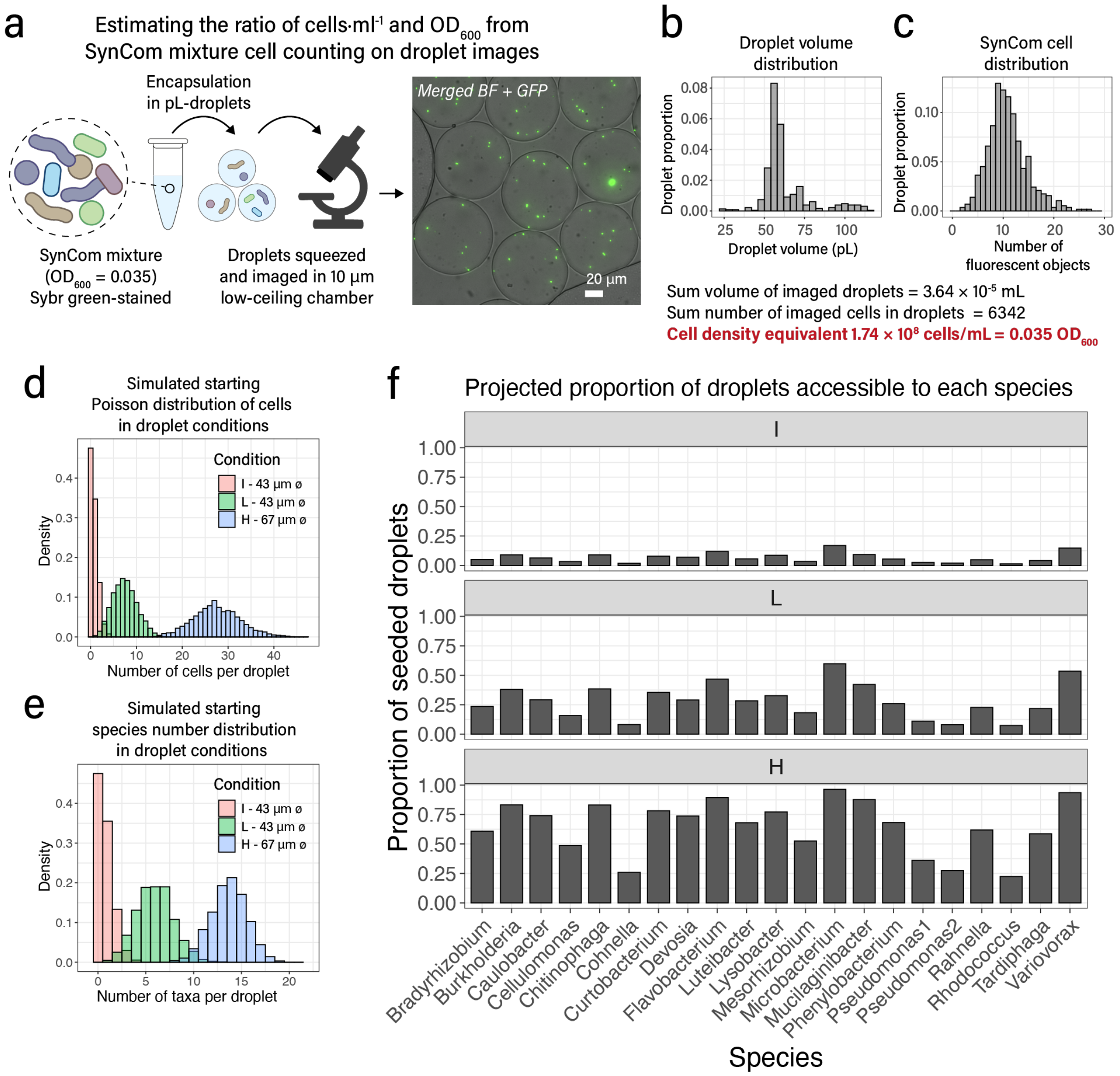


**Supplementary Figure 2: Projection of the starting species distributions in droplets**. **a** Estimated starting distribution of SynCom cells in droplets from imaging of a SynCom mixture prepared at $OD_{600}$ = 0.035, cells prestained with SYBR green I, and encapsulated in droplets (image example from droplets injected inside a low-ceiling 10 µm chamber to improve the focusing of encapsulated cells. **b**) Distribution of imaged droplet volumes and **c**) of cells per droplet, used to infer the initial cell density in the starting SynCom mixture at an $OD_{600}$ = 0.035. **d**) Projected Poisson cell number distributions in the I, L and H species level order conditions, based on the average droplet sizes measured (see Supplementary Fig. 8) and starting SynCom densities ($OD_{600}$ = 0.0035 for I, and 0.035 for L and H). **e**) Random species attribution to each cell based on the previously estimated starting proportions (Supplementary Fig. 1) to deduce the distribution of species starting numbers in droplets. **f**) Fraction of droplets in which species were present, to approximate the fraction of the accessible habitat to each species.

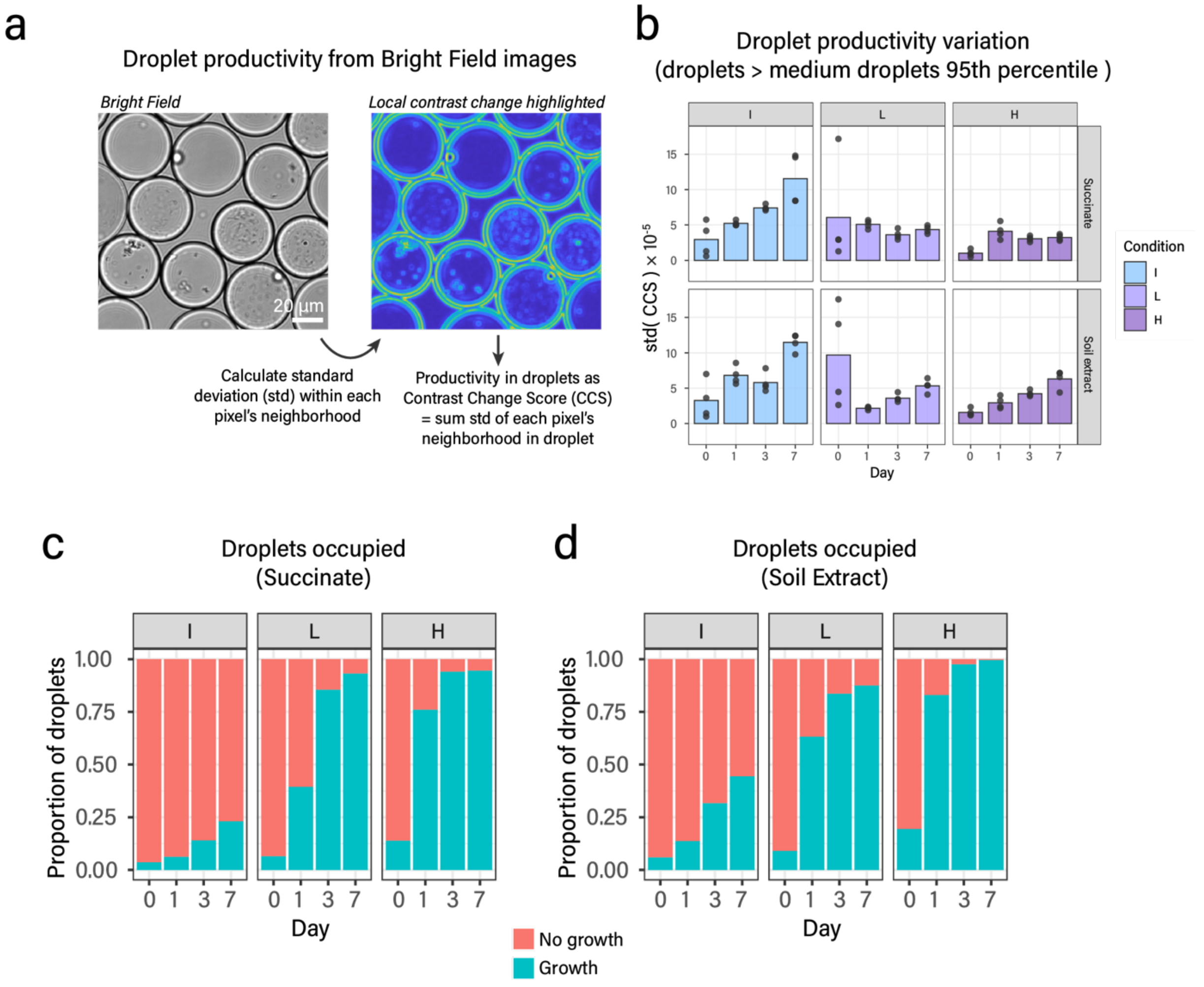


**Supplementary Figure 3: Droplet image analysis. a**) Bright-field image processing to quantify the biomass based on a local score of contrast variation (Contrast Change Score, see methods for details). The boundaries of objects with sharp edges, such as droplets and bacterial cell objects, exhibit a high local standard deviation in the gray scale values (right panel picture). **b**) Variation in droplet productivities for which the CCS is above the 95$^{th}$ percentile of imaged empty medium droplets, for both succinate and soil extract. Mean proportion (n = 4) of droplets exhibiting a CCS above the 95$^{th}$ percentile of medium droplets CCS **c**) for the succinate and **d**) for the SE condition. The gradual increase of the droplet fraction exhibiting growth illustrates the heterogeneity of growth across individual droplets.

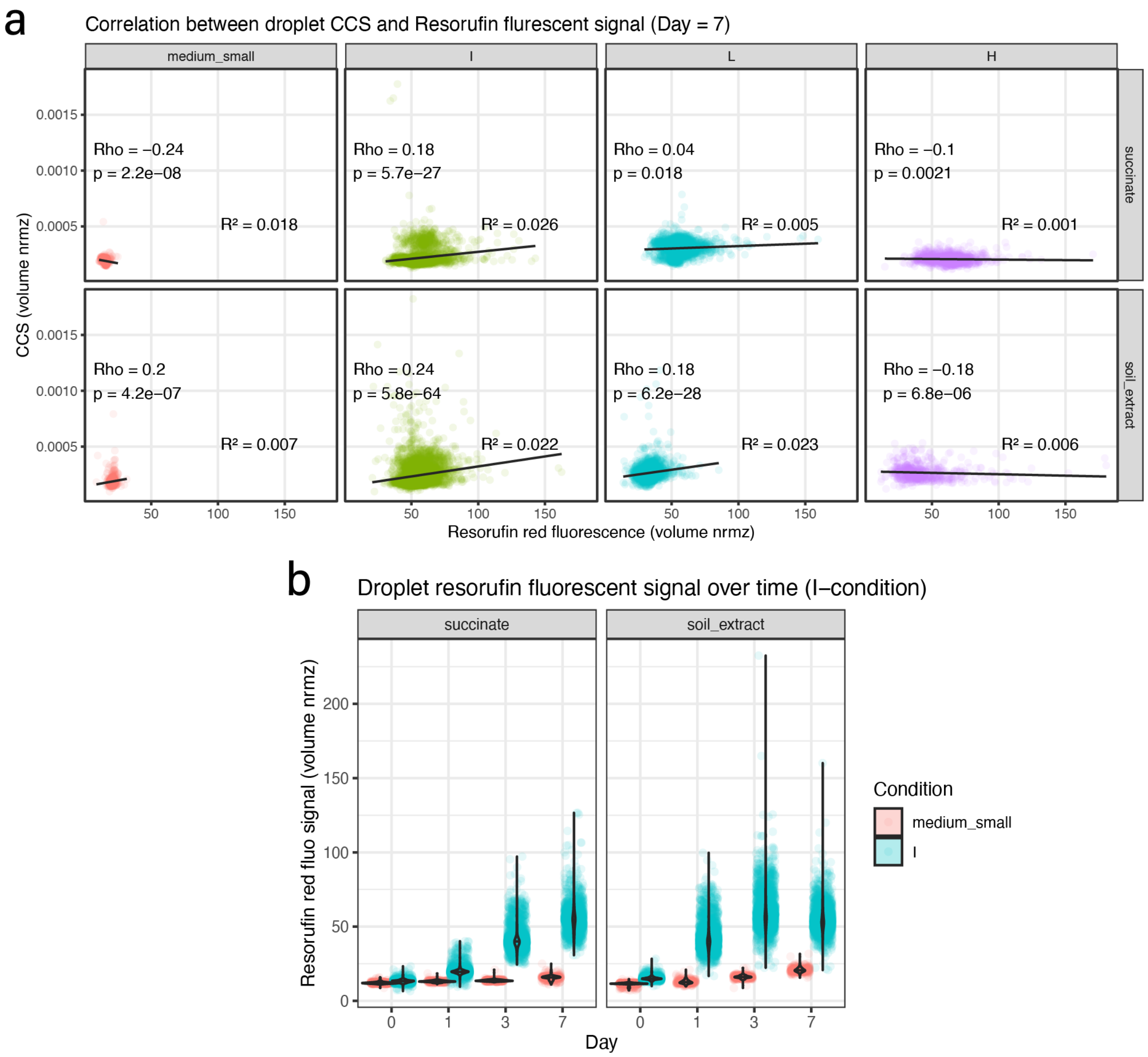


**Supplementary Figure 4: Correlation between resorufin red fluorescent signal and bright field CCS.** **a**) Scatter plots of the resorufin red fluorescent signal (volume normalized), imaged in droplets at day 7, versus the imaged CCS values estimated from the bright field channel (volume normalized). Gray lines indicate linear regression trendlines between the two measured values, and $R^2$ indicates the variation in the data points explained by the regression. Rho: Spearman correlation coefficient displayed with associated p-values. Data points from individual imaged droplets, in both succinate and soil extract. The results indicate a poor correlation between the two metrics. In particular, droplets with a low CCS, like the CCS values of medium-only droplets (*i.e.*, empty droplets), display a similar resorufin fluorescent signal as droplets with a high CCS after 7 days (panel a) and Fig. 1d), suggesting that the resorufin diffuses between droplets and cannot be used as a reliable growth reporter for single droplets. This is illustrated in **b**) with the isolated-condition droplets (I-state), where the red fluorescent signal increases for all droplets above the level of fluorescence medium-only droplets (*i.e.,* no bacteria).

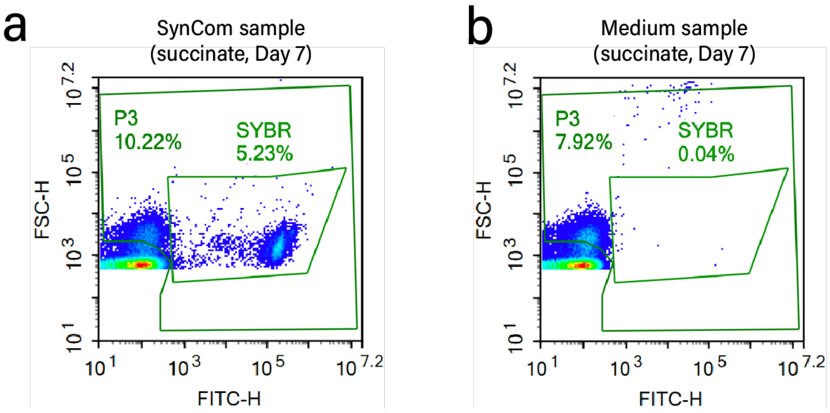


**Supplementary Figure 5: Gating strategy to identify SynCom cells among flow-cytometry events. (a)** Example scatter plots generated upon flow-cytometry counting of SYBR-stained SynCom cells showing FSC-H (thresholded to above a value of 400) and FITC-H (green fluorescence) of recorded events. The SYBR gate is drawn to count objects with a higher green fluorescence than the background SYBR-stained medium, as shown in **(b)**.

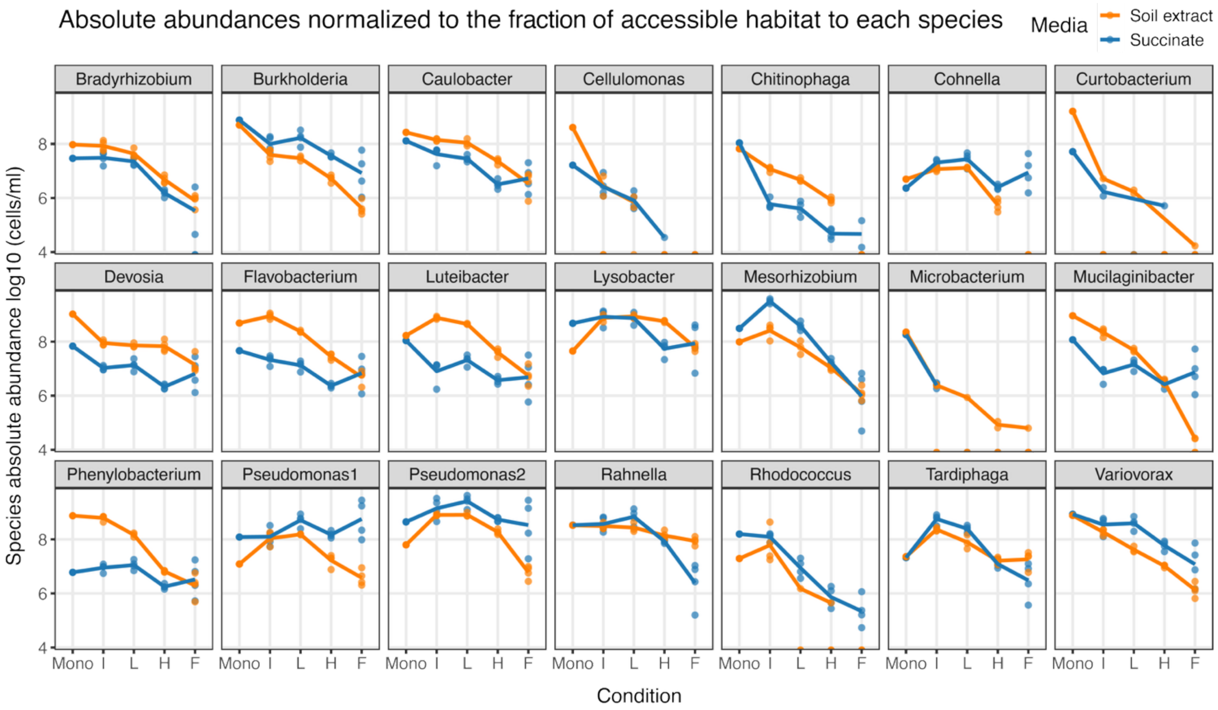


**Supplementary Figure 6: Absolute species abundances after 7 days of SynCom growth.** Absolute abundances were calculated by multiplying relative abundances (Fig. 2, from 16S rRNA gene amplicon sequencing) by the total community sizes measured by flow cytometry. 'Mono' indicates the abundances in liquid monocultures (curves presented in Supplementary Fig. 7). Differences between the abundances measured in liquid monocultures and the ones inferred from the isolation droplet scenario (I) may be due to either a technical bias or an impact of different cultivation settings (96-well plate versus droplet emulsions in Eppendorf tubes).

104

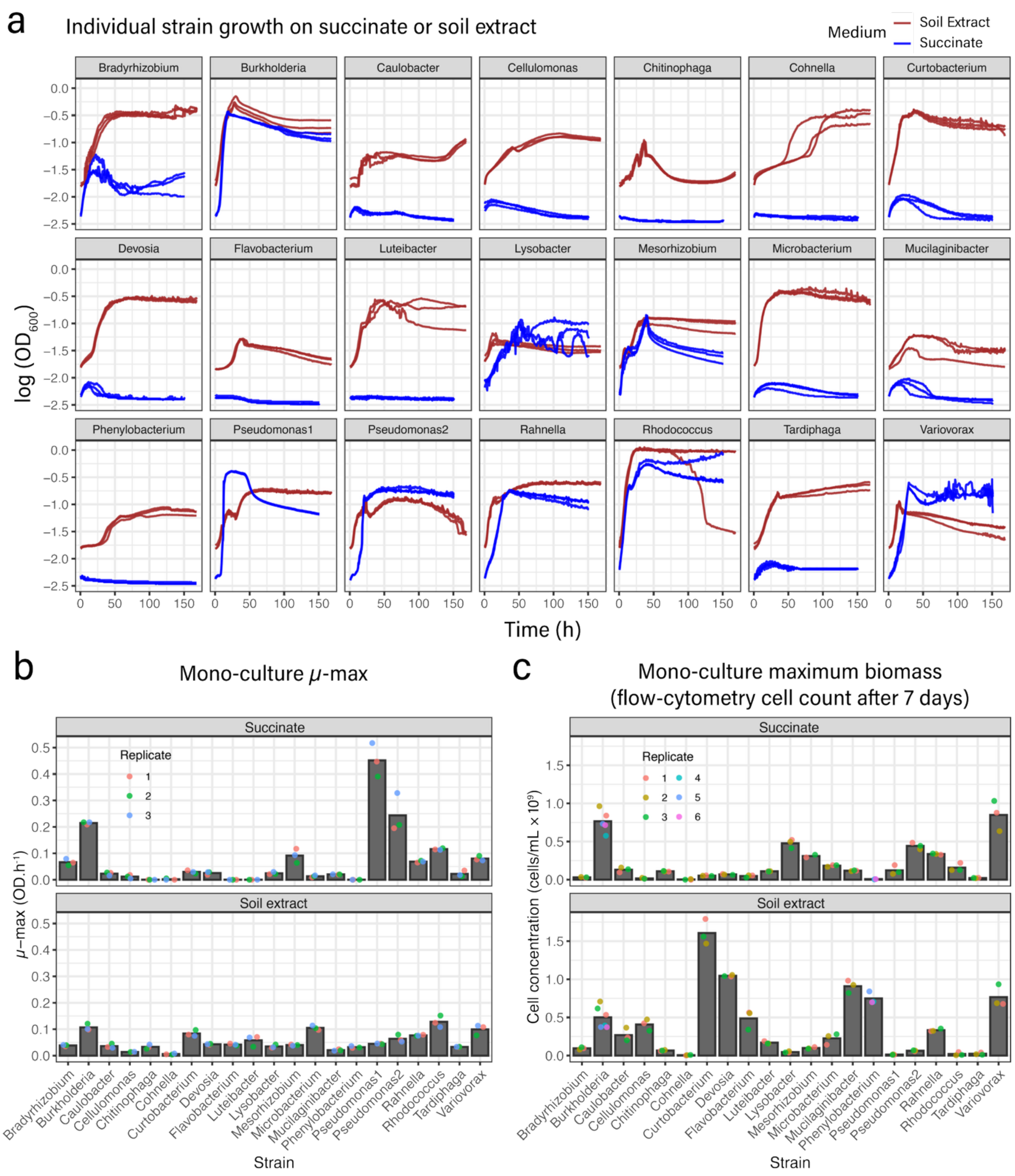


105

106 **Supplementary Figure 7: Monoculture bulk liquid growth kinetics.** a) Growth curves from LN-transformed $OD_{600}$ signals for each SynCom member monoculture on minimal medium with succinate or soil extract (n = 3) over 7 days. b) Species growth rates derived from the LN-transformed growth curves presented in a). Bars indicate the mean growth rate, and dots correspond to individual triplicate values. c) Species maximum observed number of cells per mL. Bars indicate means from triplicate measurements.

111

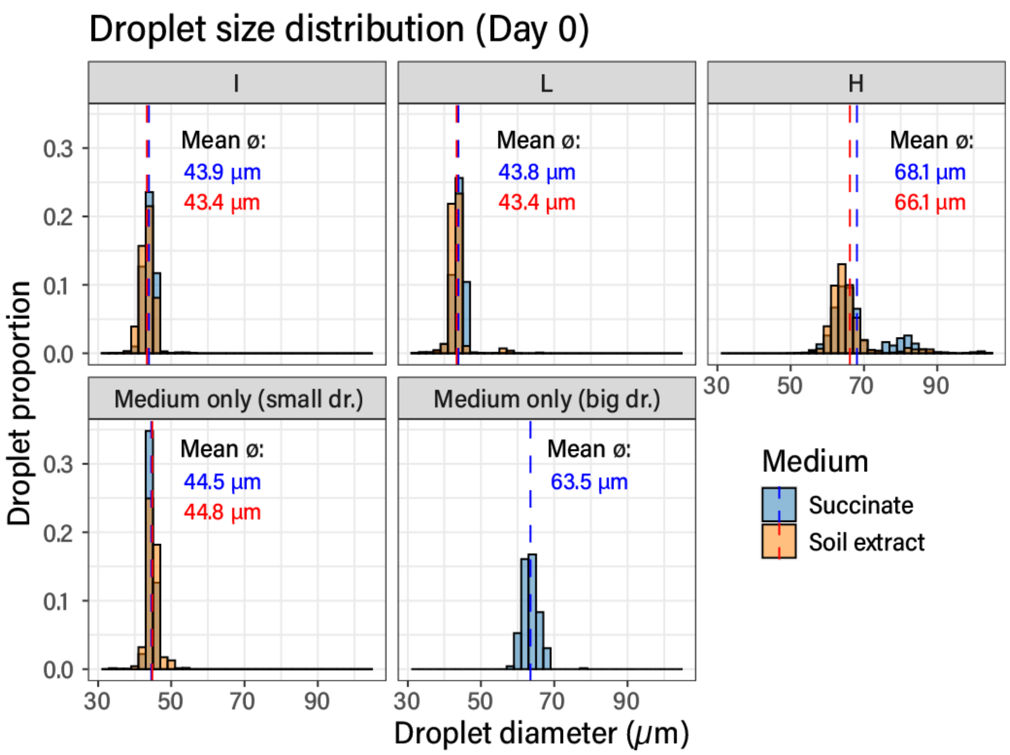


**Supplementary Figure 8: Size distribution of droplets after cell encapsulation at Day 0.** Sizes estimated from droplet microscopy images taken for each condition and medium (pooled replicates, n = 4). Dashed blue and red lines indicate the mean diameter for succinate and soil extract droplets, respectively. The significant overlap of the distributions suggests that the medium did not affect the size of the generated droplets.

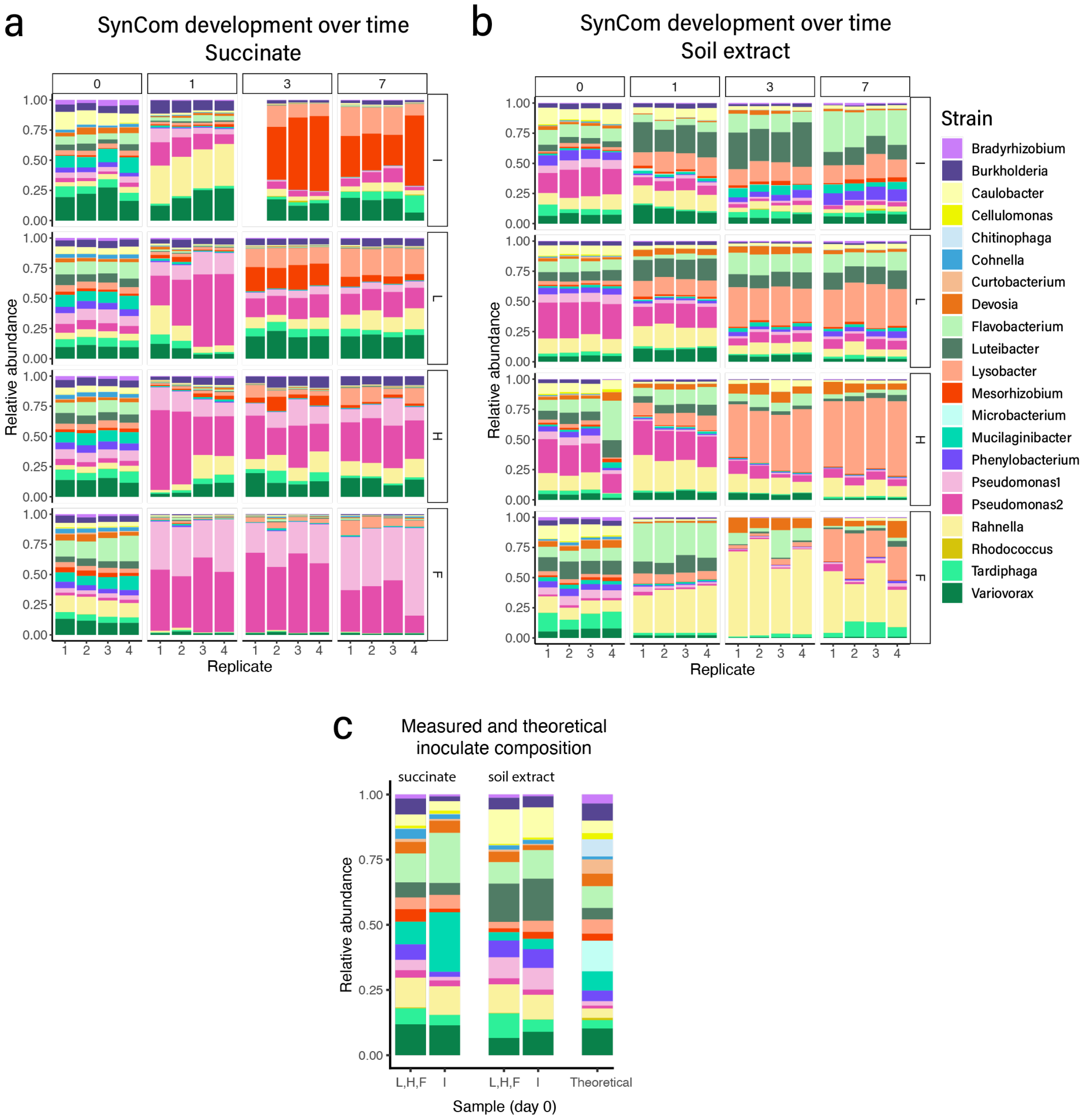


**Supplementary Figure 9: Compositional variation of the SynCom over time in the different media and conditions.** Stacked barplots of the relative abundances of the 21 SynCom members over time, as estimated from 16S rRNA gene amplicon sequencing on subsampled communities, when growing in **a**) succinate or in **b**) soil extract medium. **c**) Measured and predicted inoculant composition (as in Supplementary Fig. 1). The difference between the predicted and measured inoculant composition suggests potential biases in the DNA extraction efficiency between species.

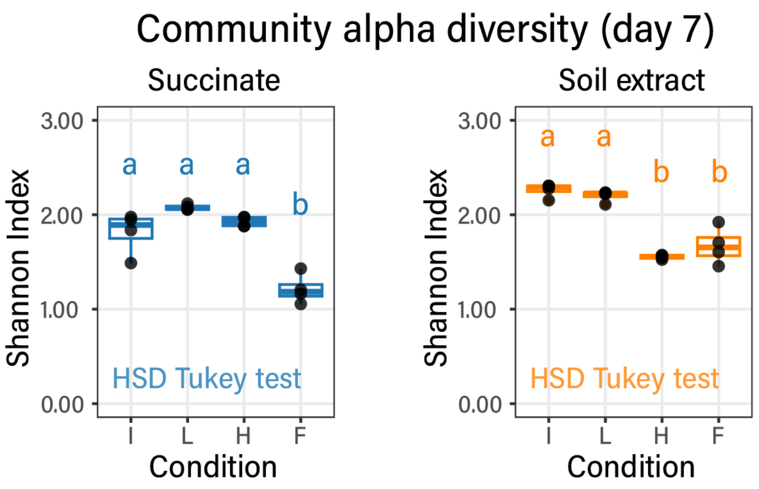


**Supplementary Figure 10: SynCom alpha diversity** on succinate and soil extract grown communities (Shannon index) after 7 days of growth for the different conditions (I, L, H, and F, as previously). Letters indicate statistically similar diversity between conditions, from HSD Tukey testing at $p < 0.05$.

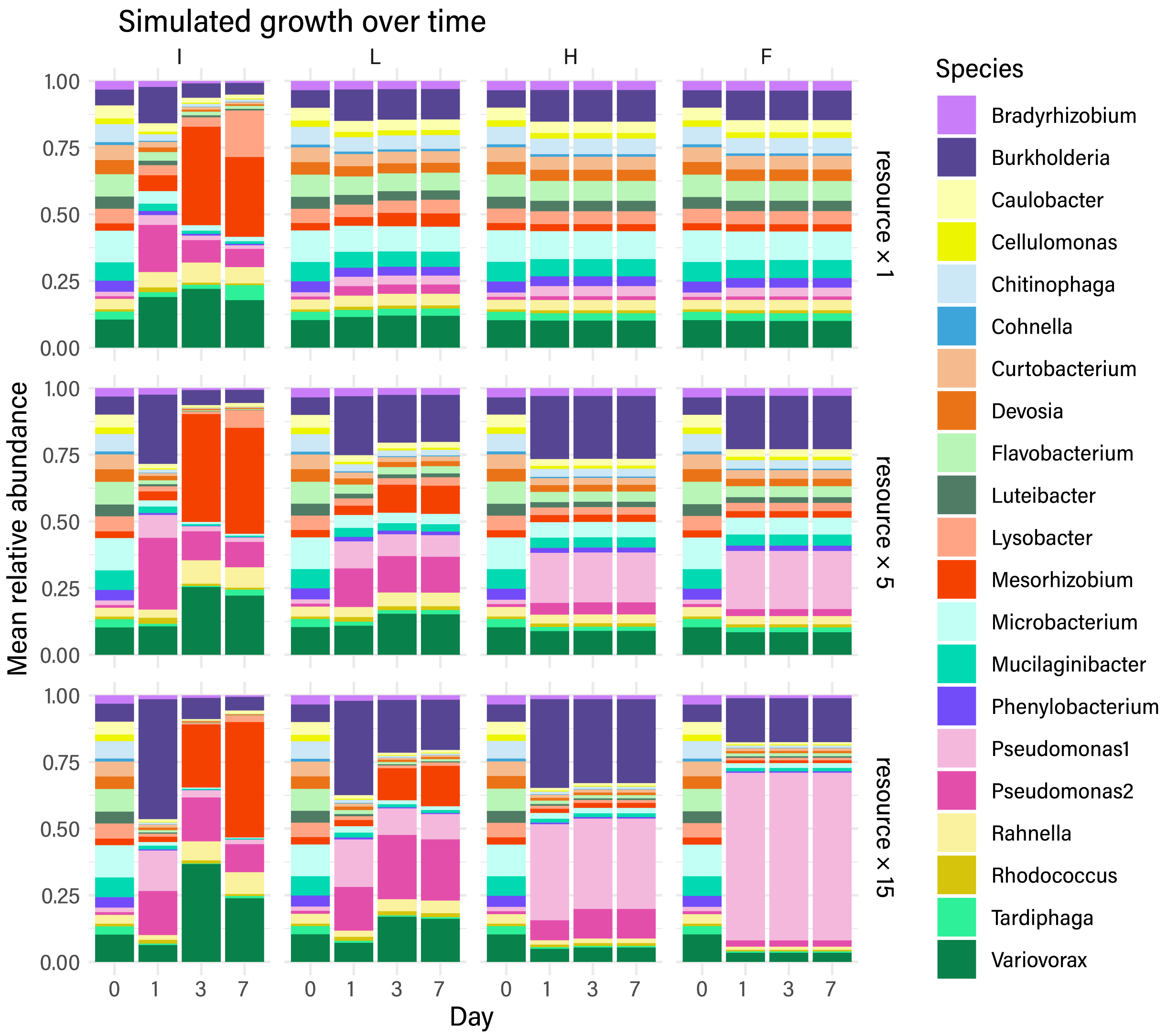


**Supplementary Figure 11: Simulated compositional variation of the SynCom over time on succinate.** Stacked barplots of the relative abundances of the 21 SynCom members over time derived from a Monod-growth kinetic model on succinate. Each row corresponds to a different iteration of the simulation, where the starting resource concentration (succinate) was untouched (resource × 1) or increased by five or fifteen times, but the stochastic starting distribution of species cells remained the same in each case.

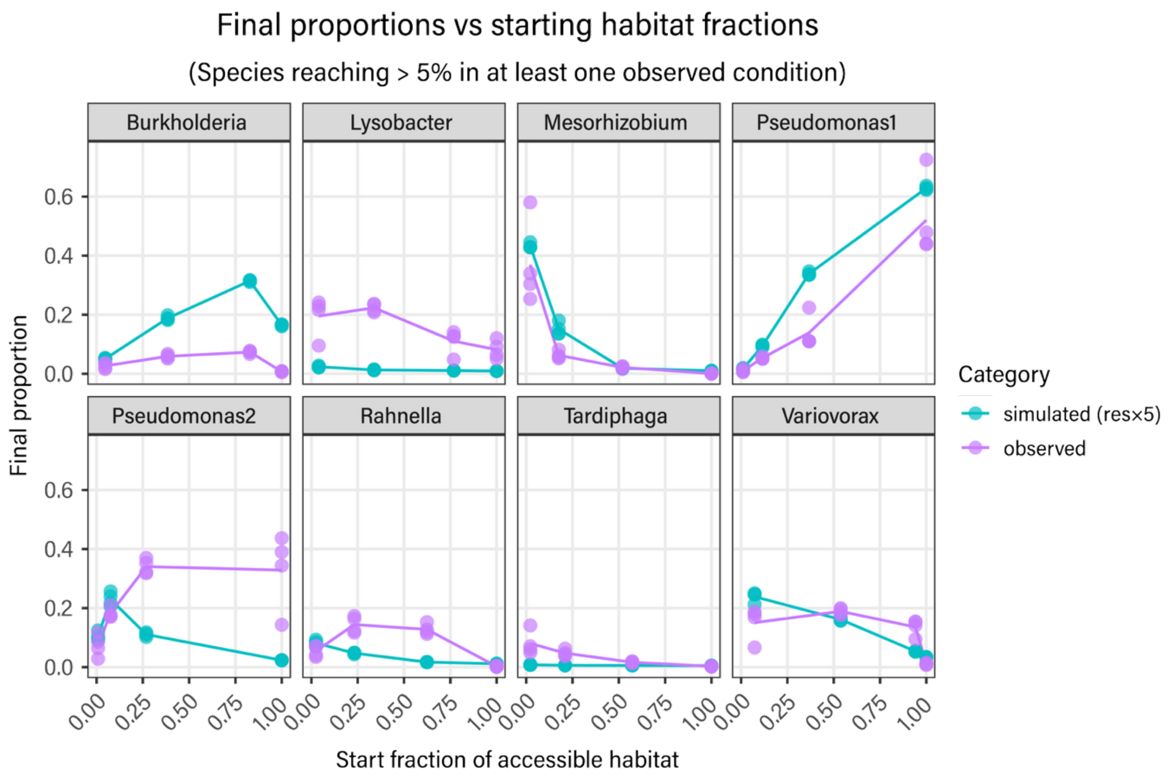


**Supplementary Figure 12: Observed and simulated species proportions** on succinate after 7 days of growth versus the accessible habitat fraction for each species calculated for each fragmentation state (see Supplementary Fig. 2). Only species reaching a minimum observed proportion of 5% of the community are shown. Each group of dots corresponds to the proportion attained in one of the conditions. Lines connect the mean proportions across accessible habitat fractions.
